# A curated list of genes that control elemental accumulation in plants

**DOI:** 10.1101/456384

**Authors:** Lauren Whitt, Felipe Klein Ricachenevsky, Greg Ziegler, Stephan Clemens, Elsbeth Walker, Frans JM Maathuis, Philip Kear, Ivan Baxter

## Abstract

Understanding the mechanisms underlying plants’ adaptation to their environment will require knowledge of the genes and alleles underlying elemental composition. Modern genetics is capable of quickly, and cheaply indicating which regions of DNA are associated with particular phenotypes in question, but most genes remain poorly annotated, hindering the identification of candidate genes. To help identify candidate genes underlying elemental accumulations, we have created the known ionome gene (KIG) list: a curated collection of genes experimentally shown to change uptake, accumulation, and distribution of elements. We have also created an automated computational pipeline to generate lists of KIG orthologs in other plant species using the PhytoMine database. The current version of KIG consists of 176 known genes covering 5 species, 23 elements and their 1588 orthologs in 10 species. Analysis of the known genes demonstrated that most were identified in the model plant *Arabidopsis thaliana*, and that transporter coding genes and genes altering the accumulation of iron and zinc are overrepresented in the current list.

## Introduction

Understanding the complex relationships that determine plant adaptation will require detailed knowledge of the action of individual genes, the environment and their interactions. One of the fundamental processes that plants must accomplish is to manage the uptake, distribution and storage of elements from the environment. Many different physiological, chemical, biochemical and cell biology processes are involved in moving elements, implicating thousands of genes in every plant species. Modern genetic techniques have made it easy and inexpensive to identify hundreds to thousands of loci for traits, such as, the elemental composition (or ionome) of plant tissues. However, moving from loci to genes is still difficult as the number of possible candidates is often extremely large and the ability of researchers to identify a candidate gene from its functional annotations is limited by our current knowledge and inherent biases about what is worth studying (Stoeger et al. 2018; I. Baxter 2020).

The most obvious candidates for genes affecting the ionome in a species are orthologs of genes that have been shown to affect elemental accumulation in another species. Indeed, there are multiple examples of orthologs affecting elemental accumulation in distantly related species, such as *Arabidopsis thaliana* and rice (*Oryza sativa*), including Na^+^ transporters from the HKT family (Z.-H. Ren et al. 2005; I. Baxter et al. 2010); the heavy metal transporters AtHMA3 and OsHMA3 (Chao et al. 2012; Jiali Yan et al. 2016); E3 ubiquitin ligase BRUTUS and OsHRZs that regulate degradation of iron uptake factors (Selote et al. 2015; Hindt et al. 2017; T. Kobayashi et al. 2013) and the K^+^ channel AKT1 (Lagarde et al. 1996; Ahmad, Mian, and Maathuis 2016). To our knowledge, no comprehensive list of genes known to affect elemental accumulation in plants exists. To ameliorate this deficiency, we sought to create a curated list of genes based on peer reviewed literature along with a pipeline to identify orthologs of the genes in any plant species and a method for continuously updating the list. Here we present version 1.0 of the known ionome gene (KIG) list.

## Materials and Methods

The list includes all functionally characterized genes from the literature that are linked to changes in the ionome. Criteria for inclusion in the primary KIG list were as follows:

1. The function or levels of the gene are unambiguously altered (i.e. a confirmed knockout, knockdown or over expressor). For double mutants, both genes are listed.
2. The levels of at least one element are significantly altered in a plant tissue.
3. Publication in the form of a peer reviewed manuscript.

Note that our definition excludes genes that are linked to metal tolerance or sensitivity but do not alter the ionome, or genes where the levels of the transcript are correlated with elemental accumulation. In order to identify the KIG genes, we created a Google survey that was distributed to members of the Ionomicshub research coordination network (NSF DBI-0953433), as well as advertising on Twitter and in oral presentations by the authors. We asked submitters to provide the species, gene name (or names where alleles of two genes were required for a phenotype), gene ID(s), tissue(s), element(s) altered and a DOI link for the primary literature support. Subsequently, authors FKR and LW did an extensive literature search.

### Creating the inferred orthologs list

The known ionome gene list contains known genes from the primary list and their orthologous genes inferred by InParanoid (v4.1) pairwise species comparisons (Remm, Storm, and Sonnhammer 2001). The InParanoid files were downloaded from Phytozome for each organism-to-organism combination of species in the primary list, plus *Glycine max, Sorghum bicolor, Setaria italica, Setaria viridis* and *Populus trichocarpa*. Orthologs of the primary genes were labeled as “inferred” genes. If a primary gene was also found as an ortholog to a primary gene in another species, the status was changed to “Primary/Inferred” in both species. It is important to note that only primary genes can infer genes; inferred genes cannot infer other genes. The pipeline for transforming the primary list into the known ionomics gene list can be found at https://github.com/baxterlab/KIG.

### Gene Enrichment analysis

Overrepresentation analysis (released 07-11-2019) was performed on the primary and inferred genes in *A. thaliana* using the GO Consortium’s web-based GO Enrichment Analysis tool powered by the PANTHER (v14) classification system tool (Ashburner et al. 2000; The Gene Ontology Consortium 2017; Mi et al. 2017). We restricted overrepresentation analysis to *A. thaliana* because of its dominance in the KIG list and our lack of confidence in the functional annotation of the other species in the list. An analysis performed by Wimalanathan et al. (2018) found that maize gene annotations in databases like Gramene and Phytozome lacked GO annotations outside of automatically assigned, electronic annotations (IEA). IEA annotations are not curated and have the least amount of support out of all the evidence codes (Harris et al. 2004). *A. thaliana* annotations come from a variety of evidence types, showing a higher degree of curation compared to maize (Wimalanathan et al. 2018). The whole-genome *Arabidopsis thaliana* gene list from the PANTHER database was used as the reference list.

We tested both the PANTHER GO-slim and the GO complete datasets for biological processes, molecular function and cellular component. GO-Slim datasets contain a selected subset of terms that give a broad summary of the gene list, whereas the complete dataset contains all the terms returned for a more detailed analysis. The enriched terms (fold enrichment > 1 and with a false discovery rate <0.05) from the complete dataset were sorted into five specific categories relating to the ionome based annotation terms:

1. Ion homeostasis - terms include homeostasis, stress, detoxification, regulation of an ion
2. Ion transport - terms specifically state transport, export, import or localization of ion(s). Does not include hydrogen ion transport
3. Metal ion chelation - terms relating to phytochelatins, other chemical reactions or pathways of metal chelator synthesis
4. Response to ions - vaguely states a response to ions, but does not have any parent annotation terms that offer any more clarification (ie. stress response). Broadly this is referring to any change to the state or activity of cell secretion, expression, movement, or enzyme production (Carbon et al. 2009)
5. Other transport - annotation stating the transfer of anything that is not an ion (glucose, peptides, etc.)

Genes may belong to more than one category, but if they belong to a parent and child term in the same category, they are only counted once.

## Results

The current primary list (v1.0) consists of 176 genes from *A. thaliana, O. sativa, Medicago truncatula, Triticum aestivum* and *Zea mays* with the majority coming from *A. thaliana* and *O. sativa* (Table 1)(Figure 1).

**Table 1.**
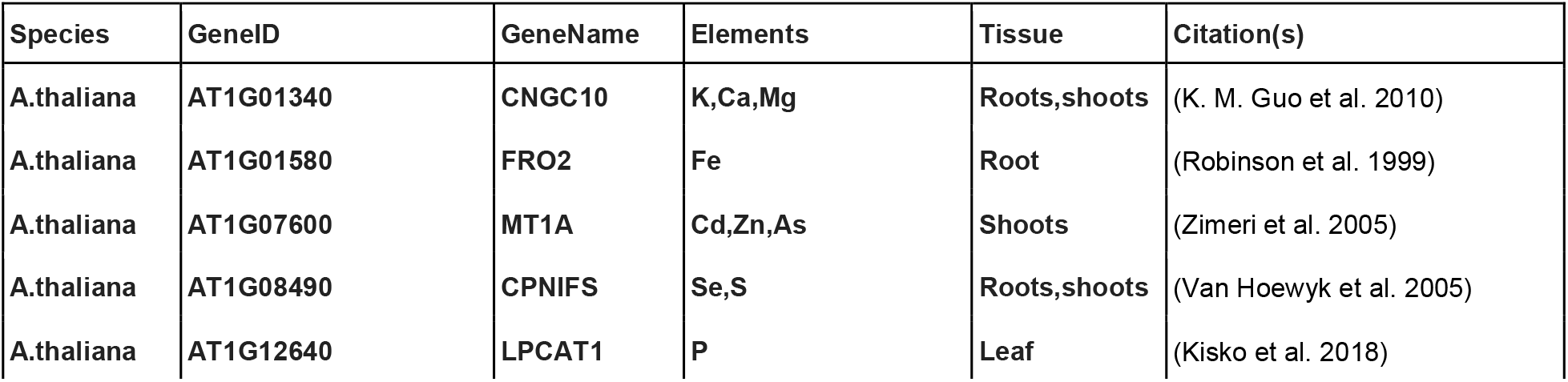
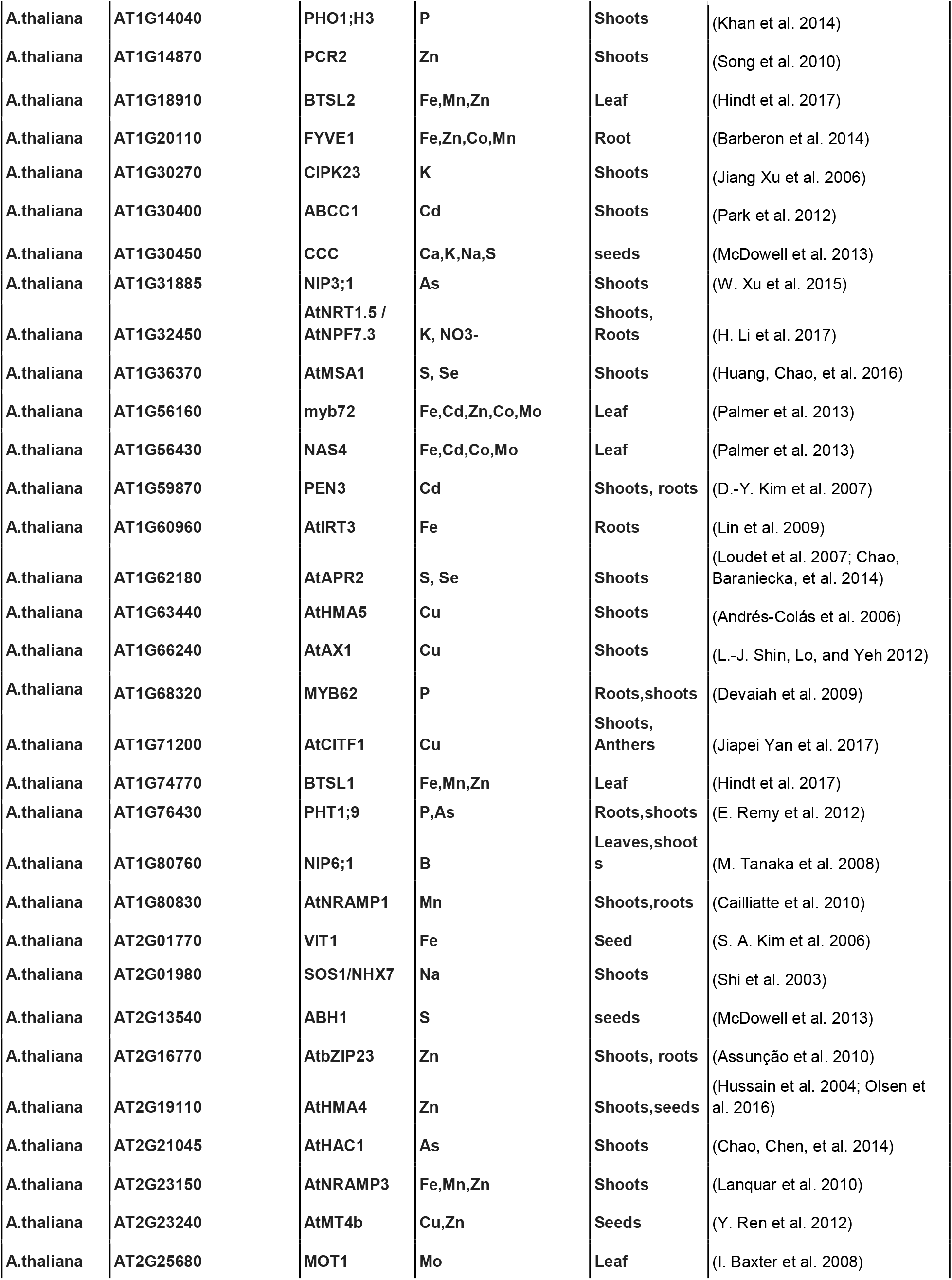
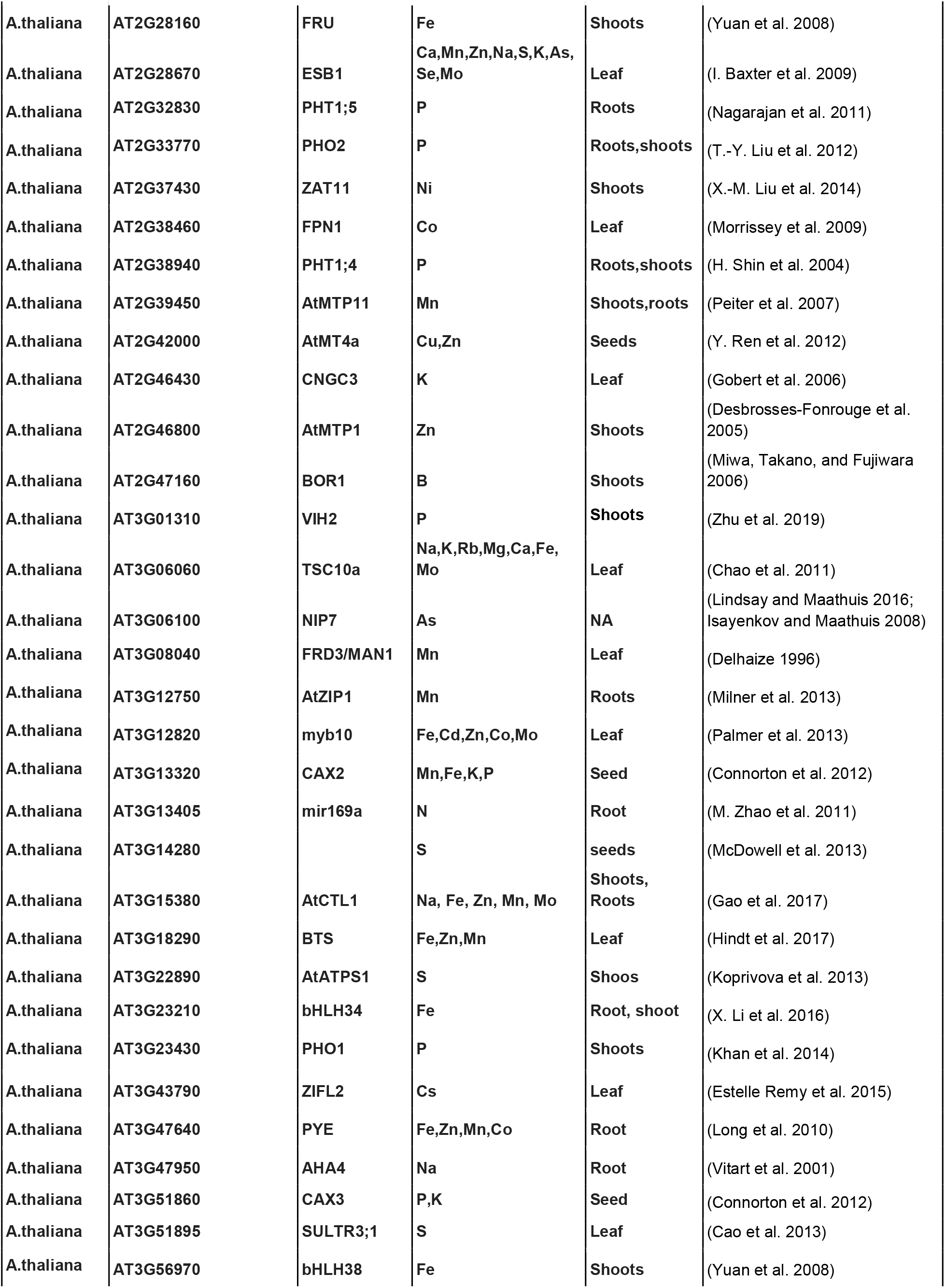
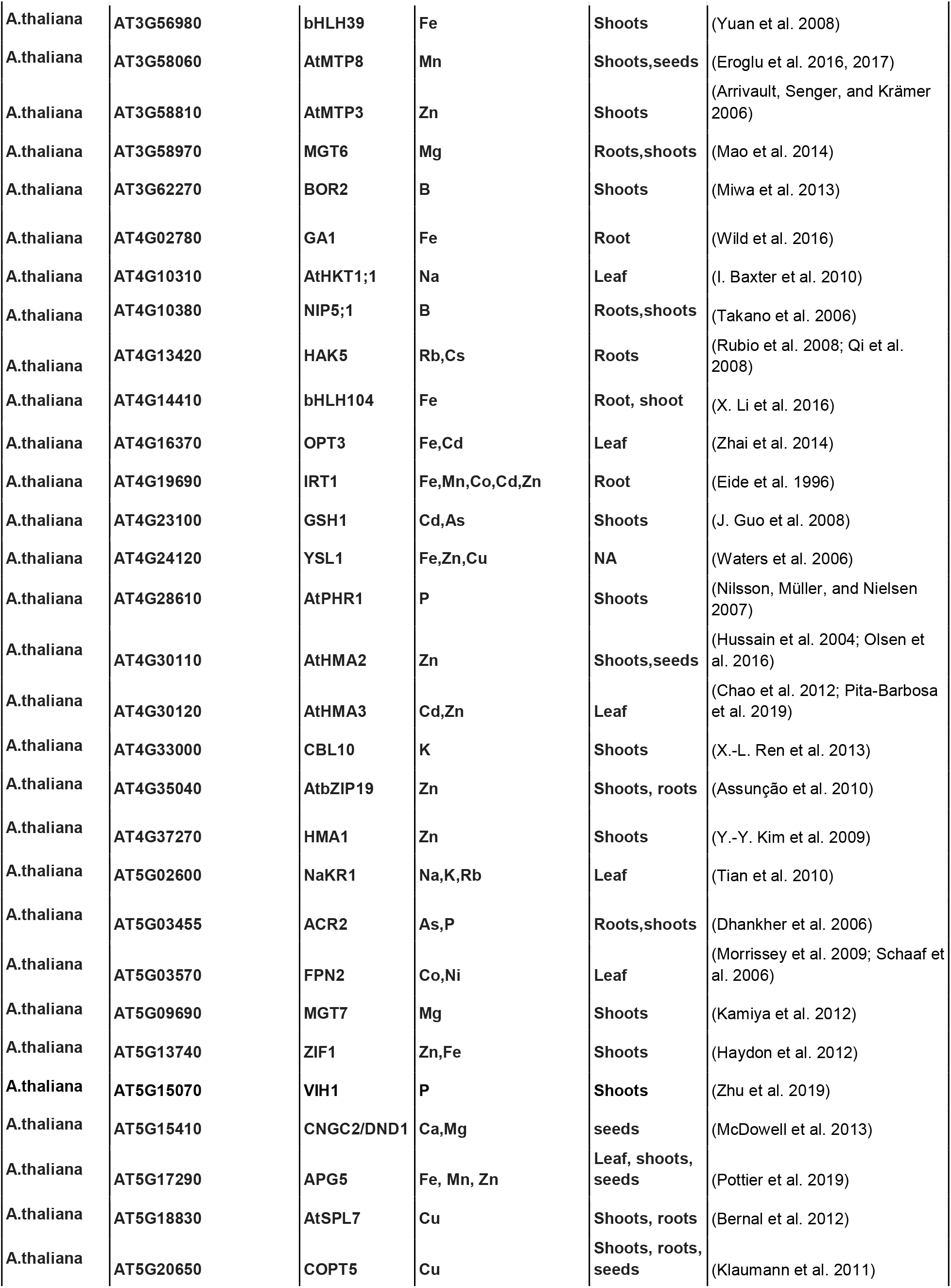
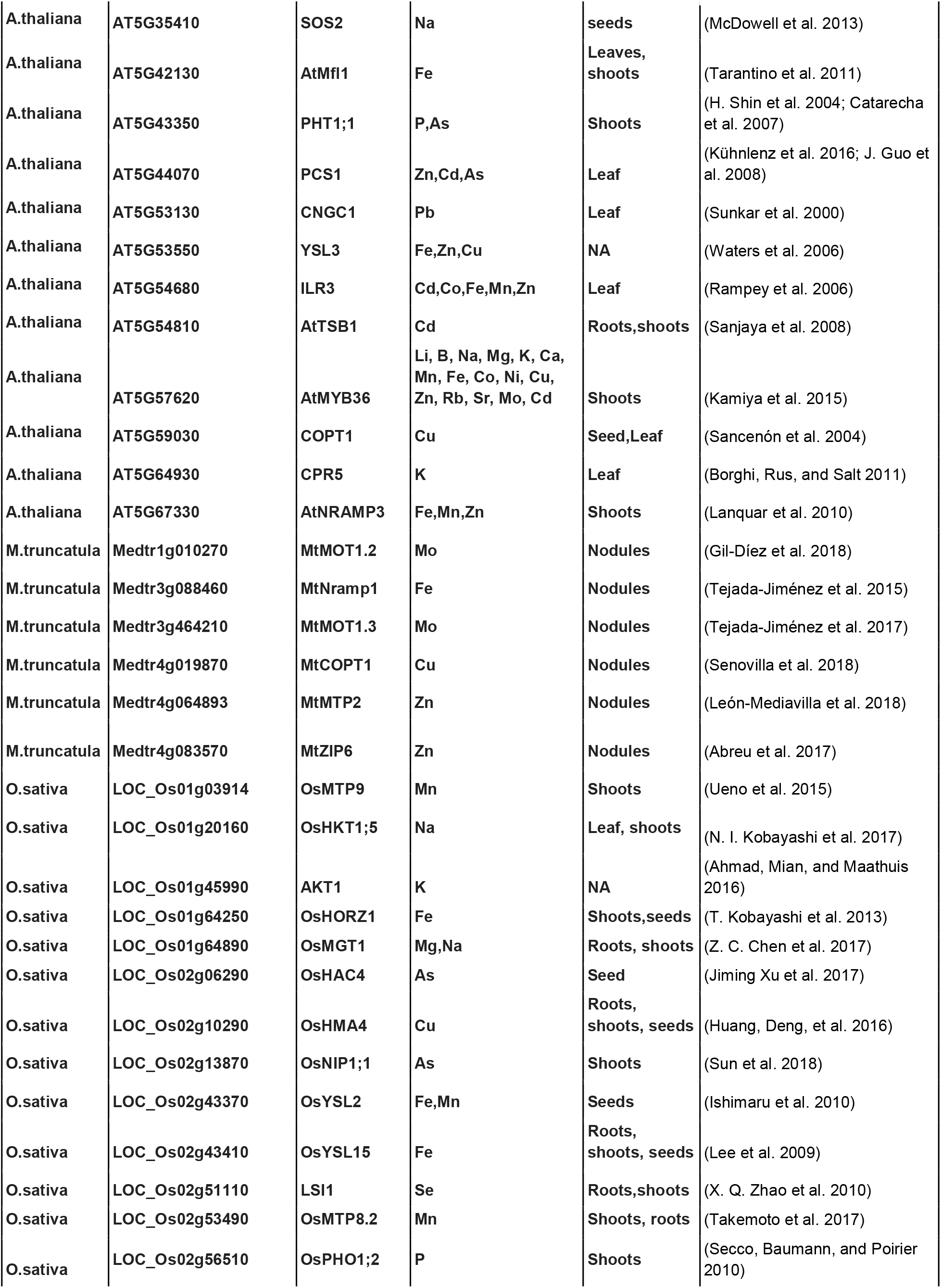
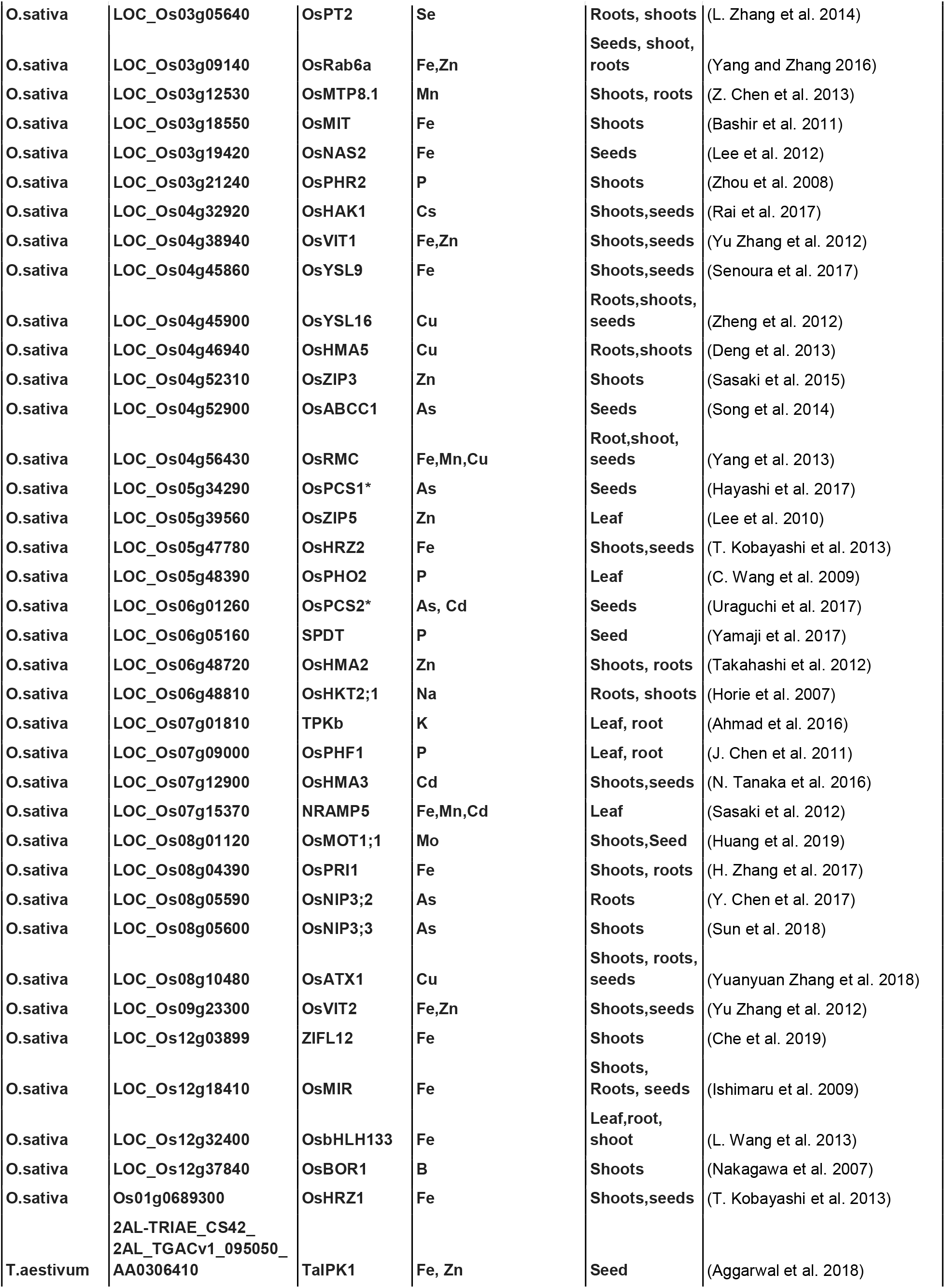
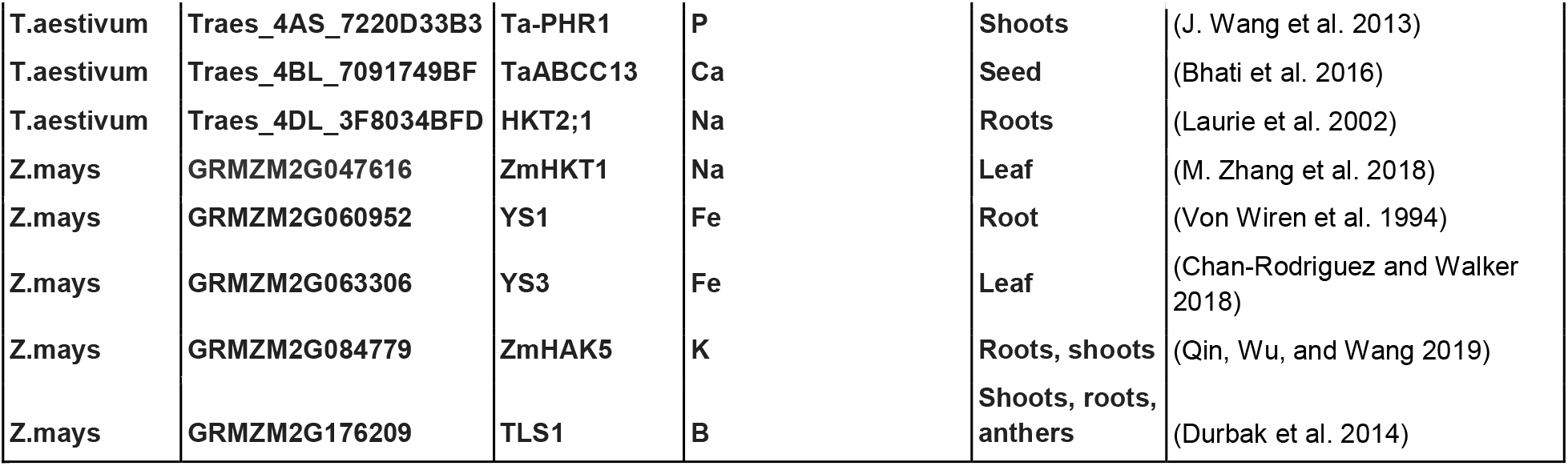
Primary known ionome genes.

**Figure 1.**
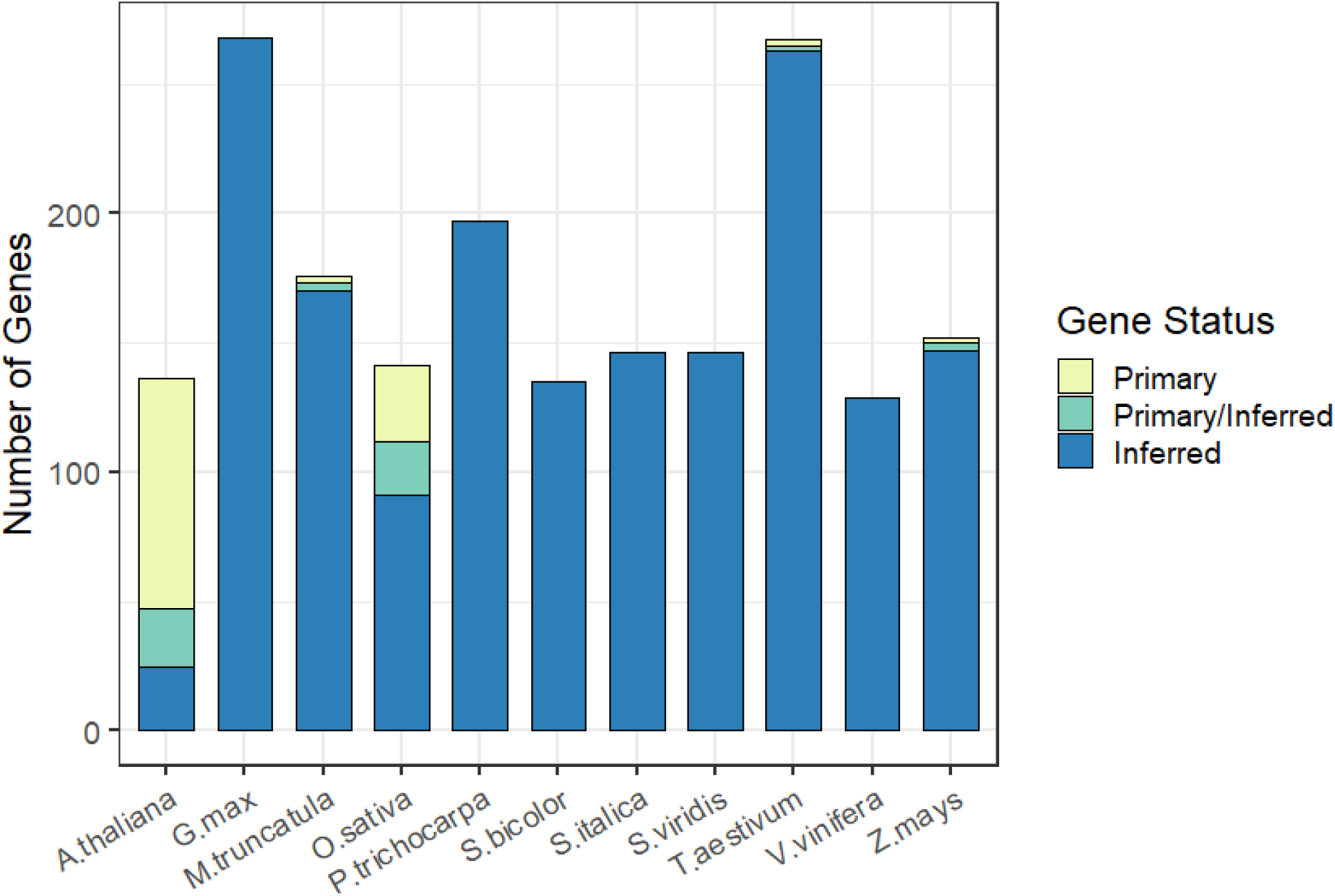
Number of genes for each species that are primary, inferred from other primary genes in other species, or both.

Most primary genes have orthologs in other species. Less than 10% of primary genes in *A. thaliana*, 12% in *O.sativa* and one of the four primary genes in wheat (*T. aestivum*) lack orthologs (Table 2). *G. max, P.trichocarpa, S. bicolor, S. italica*, and *S. viridis* currently contain only inferred genes (Table 2, Figure 1).

**Table 2.** Break down of primary/inferred genes in each species.

| **Table 2. Break down of primary/inferred genes in each species.** |  |  |  |  |  |
| --- | --- | --- | --- | --- | --- |
| **Species** | **Total Genes** | **Primary Genes** | **Primary/Inferred Genes** | **Inferred Genes** | **Primary & Primary/Inferred Genes without orthologs** |
| A.thaliana | 136 | 65.44% | 16.18% | 18.38% | 9.91% |
| O.sativa | 141 | 20.57% | 14.89% | 64.54% | 12.00% |
| M.truncatula | 176 | 1.70% | 1.70% | 96.59% | 0.00% |
| T.aestivum | 267 | 0.75% | 0.75% | 98.50% | 25.00% |
| Z.mays | 152 | 1.32% | 1.97% | 96.71% | 0.00% |
| G.max | 268 | 0.00% | 0.00% | 100.00% | 0.00% |
| P.trichocarpa | 197 | 0.00% | 0.00% | 100.00% | 0.00% |
| S.bicolor | 135 | 0.00% | 0.00% | 100.00% | 0.00% |
| S.italica | 146 | 0.00% | 0.00% | 100.00% | 0.00% |
| S.viridis | 146 | 0.00% | 0.00% | 100.00% | 0.00% |

The YSL genes in *A. thaliana* and *O.sativa* are an example that provides evidence for the validity of the KIG list pipeline: AtYSL3, OsYSL9 and OsYSL16 were listed in their respective species as primary genes (Table 1) and after the ortholog search were annotated as primary/inferred genes, referencing each other (STable1). AtYSL2 in *A. thaliana*, was not listed as primary gene, but was inferred through its rice orthologs OsYSL9 and OsYSL16. Additionally, AtYSL1 in *A. thaliana* is not a paralog of AtYSL3 or an ortholog of OsYSL9 and OsYSL16 according to PhytoMine’s InParanoid results, and is not listed as an ortholog to either of the *O. sativa* YSL genes in the KIG list. Other examples include AtVIT1 and OsVIT1/OsVIT2 (S. A. Kim et al. 2006; Yu Zhang et al. 2012), and the vacuolar Mn transporters AtMTP8 and OsMTP8 (Eroglu et al. 2016; Z. Chen et al. 2013). Thus, we can reliably generate inferred genes and create a species-specific KIG list for any species in PhytoMine.

The primary list covers 23 elements (Figure 2) according to the reported elements from authors in the primary list, which is more elements than predicted by the GO term annotations for those genes. Some GO annotations for these genes mention only a portion of elements listed by the literature in the primary list. This may be due to GO annotation evidence codes lacking curation or biological data (IEA,ND,NAS) (Wimalanathan et al. 2018), or it may be due to alterations in one element leading to alterations in other elements (I. R. Baxter et al. 2008).

**Figure 2.**
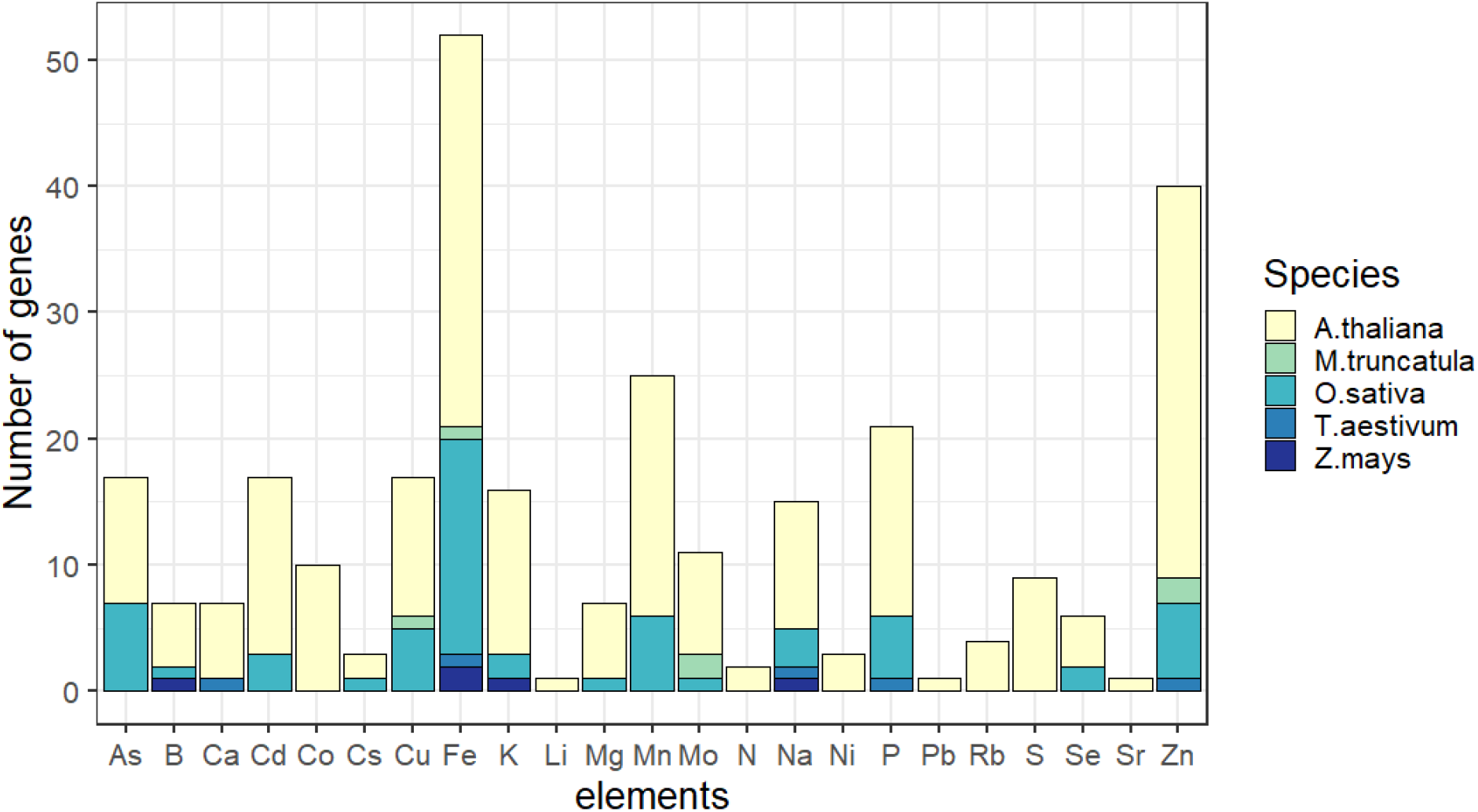
Number of primary genes from each species listing each element.

*A. thaliana* is the only species to have a primary gene listing for each element. There is a bias towards manganese, zinc and iron which have 2, 3 and 4 times more associated genes than the average 13±12 genes of other elements. Iron is the only element to contain genes from all five species in the primary list. In addition to biases towards certain elements, our primary list is also skewed towards an overrepresentation of ionome genes in above ground tissue studies (Figure 3). This is likely due to the difficulties in studying the elemental content of below ground tissues. All *M. truncatula* genes come from studies of the nodule in this model legume species.

**Figure 3.**
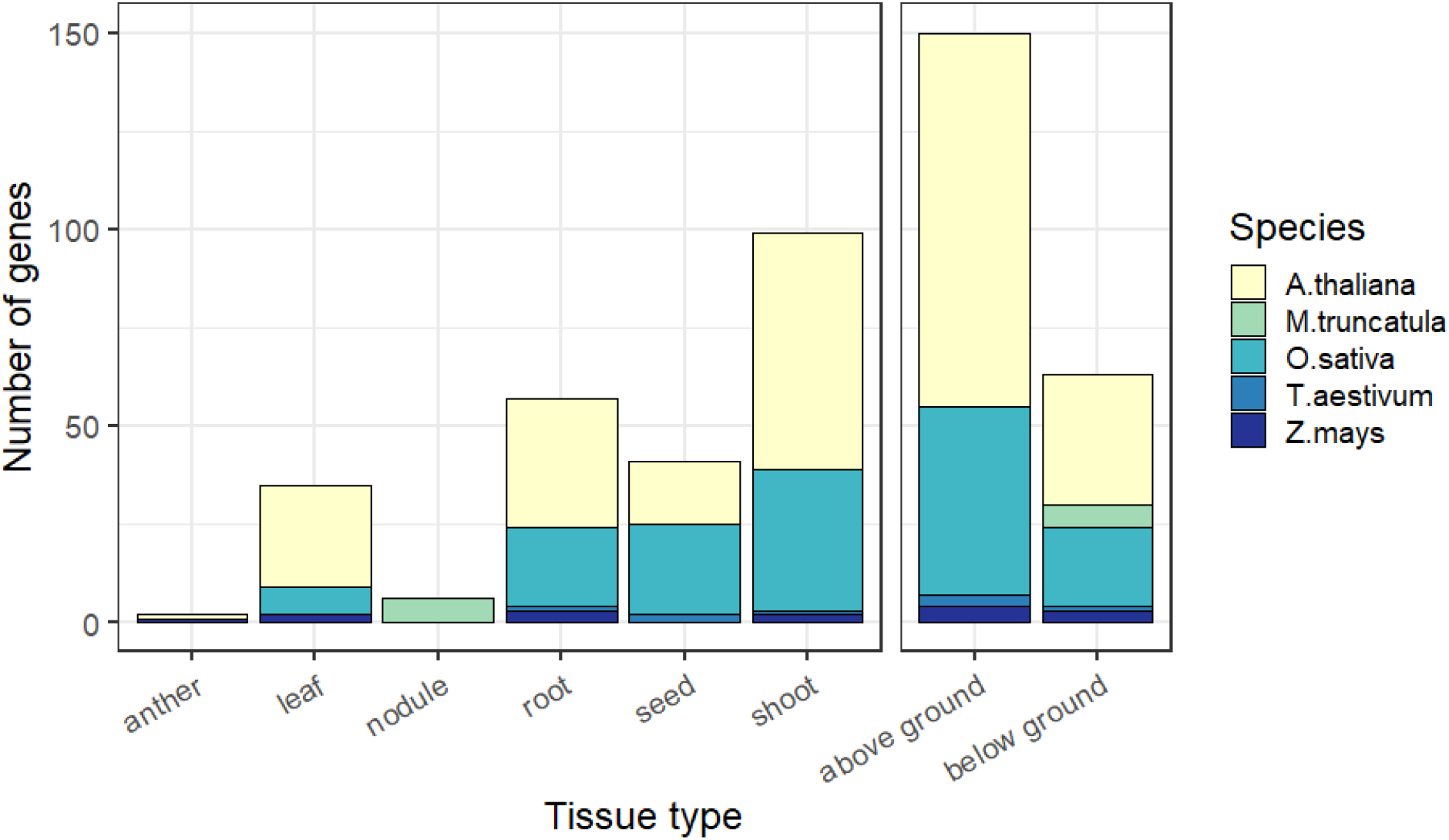
Number of primary genes each type of tissue contributes to the known ionome gene list. Above ground is a summary of anther, leaf, seed and shoot, while below ground is a summary of root and nodule.

Querying the manually curated PANTHER GO-slim biological process database (PANTHER v14.1, released 03-12-2019) and the GO complete biological process database (GO Ontology database, released 10-08-2019), with the *A. thaliana* KIG genes returned significantly (FDR < 0.05) overrepresented annotation terms related to the transport, response, and homeostasis of iron, zinc, copper and manganese ions. Additionally, the GO complete results had terms for cadmium, nickel, cobalt, sulfur, arsenic, lead, selenium, boron, magnesium, phosphorus, sodium, potassium, and calcium; covering most of the elements in the KIG list (Figure 4). Even though some genes were annotated as associated in the “other transport” of glycoside, glucose, oligopeptides, or phloem transport, the citations that have added them into our primary list show that their mutant alleles altered elemental accumulation. AtABCC1 is annotated as encoding a glycoside transporter protein, but Park et al. (2012) found overexpression of AtABCC1 increased cadmium concentrations in shoot tissue. The YSL genes and OPT3 are annotated as genes encoding oligopeptide transporters, but more specifically they are encoding predicted phloem-localized metal-nicotianamine complex and iron/cadmium transporters, respectively (Waters et al. 2006; Zhai et al. 2014). Lastly, NRT1.5/NPF7.3 is also annotated as encoding an oligopeptide transporter, but Li et al. (H. Li et al. 2017) identified it as a xylem loading potassium ion antiporter.

**Figure 4.**
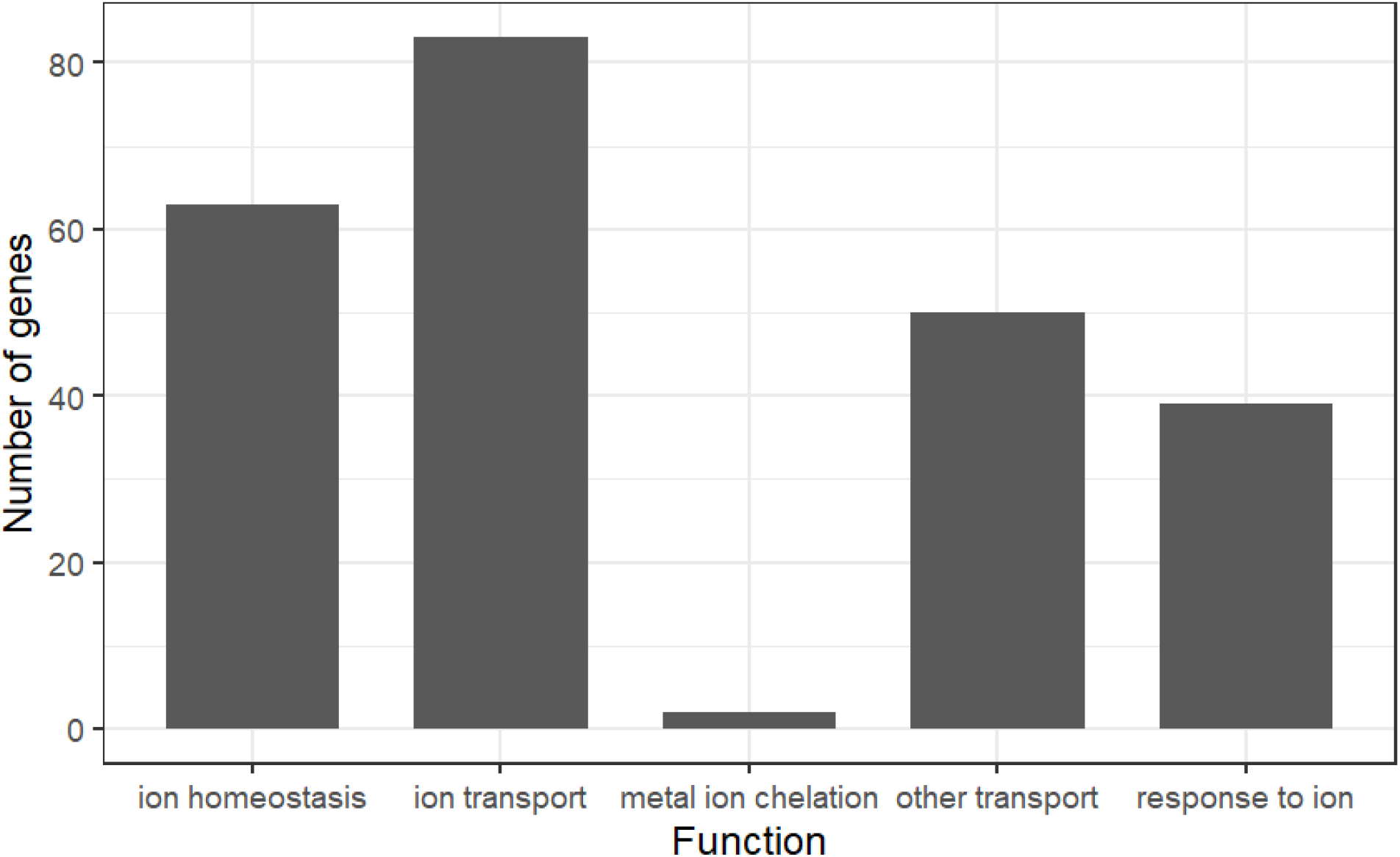
Known ionome genes relating to different terms from the GO complete biological process dataset. Ontology groups of GO Enrichment Analysis from PANTHER.

The PANTHER GO-slim molecular function annotation database found a significant overrepresentation for iron and potassium cation transmembrane transporter activity in the *A.thaliana* genes. The results using the GO complete molecular function database supported this, and additionally included terms for arsenic, cadmium, zinc, boron, manganese, phosphate, sulfur and magnesium ion transmembrane transporter activity. The GO complete molecular database also returned overrepresented terms for metal ion binding and cyclic nucleotide binding annotations. The cyclic nucleotide binding annotation genes were more specifically cyclic nucleotide ion gated channel genes (Gobert et al. 2006). The PANTHER GO-slim cell component and GO complete cell component annotation database both returned significant overrepresentation for vacuoles and the plasma membrane, both known to be critical for elemental movement and storage (Barkla and Pantoja 1996). The molecular function and cell component results are further evidence that our list is dominated by ion transporters.

To test the completeness of the KIG list, we searched PANTHER’s biological processes annotations for the number of *A. thaliana* genes encoding predicted elemental transporters. We found 618 *A.thaliana* genes predicted to encode elemental transport, and only 40 of these PANTHER genes are listed in the KIG list. We checked these results against ThaleMine (v1.10.4, updated on 06-13-2017) genes with the term “ion transport” in the gene name, description, or GO annotation and found only 358 genes, with 52 of these genes listed in the *A. thaliana* known ionome gene list. Interestingly, 219 of the genes from ThaleMine were not found in the 634 from PANTHER.

## Discussion

Here we have produced a curated list of genes known to alter the elemental composition of plant tissues. We envision several possible uses for this list:

1. Researchers can use the list to identify candidate genes in loci from QTL and GWAS experiments.
2. This list can serve as a gold standard for computational approaches.
3. The list can serve as a reading list for those interested in learning about elemental accumulation.

It is important to highlight that the inferred genes lists are not likely to be perfect predictors of the causal genes. Our use of InParanoid orthologs may exclude homologs that are likely candidates. Additionally, the reasons that some genes have been studied could be the result of human bias towards research topics (I. Baxter 2020). The list is highly enriched for 1) transporters, 2) genes that affect elemental accumulation in above ground tissues and 3) genes that affect the accumulation of Fe and Zn. Transporter genes became obvious candidates for studying plant nutrition when disruption allele collections were produced (McDowell et al. 2013). Above ground tissues are easier to study without contamination from the soil, and such studies are therefore more prevalent. Finally, while Fe and Zn are important biochemical cofactors, these elements are likely enriched in the KIG list due to their considerable interest to the community where the ionomics approach was developed. This is further illustrated in the PANTHER GO-slim databases, where Fe was the only element to have its overrepresented response, homeostasis and transport related GO terms show up in the PANTHER GO-slim biological process and molecular function databases, which are selected subsets of terms meant to broadly summarize data. Overrepresented terms related to other KIG list elements are only found in the GO complete databases. Taken together, these factors warn against forming conclusions about the nature of all elemental accumulation genes based on this limited dataset.

Most entries on this list are derived from model organisms suggesting that most of our knowledge about genes that affect elemental accumulation comes from these species. *A. thaliana* and *M. truncatula* account for 64% of the primary genes list, which is probably a lower bound for the influence of knowledge generated in model organisms. Several of the genes in crop plants were found due to being orthologs of genes in the model organisms (Ahmad, Mian, and Maathuis 2016; Jiming Xu et al. 2017), and on closer inspection of the 50 papers identifying primary genes in rice, 38 cited a gene in Arabidopsis (not necessarily the direct ortholog) as a source for why the gene was investigated.The higher quality of the GO terms in Arabidopsis when compared to other species is another reflection of this disparity of knowledge and a significant hindrance when trying to clone genes in other organisms.

## Call for more submissions

While we have done our best to ensure that the current list is useful and thorough, it is possible we are still missing genes. We ask readers who know of genes that we are missing to contribute by submitting them here: https://docs.google.com/forms/d/e/1FAIpQLSdmS_zeOlxTOLmq2wB45BuSQml1LMKtKnWSatmFRGR2Q1o0Ew/viewform?c=0&w=1 or email corresponding author. KIG lists v1.0 for each of the species can be viewed in STable1, and future updates to the list can be found at https://docs.google.com/spreadsheets/d/1XI2l1vtVJiHrlXLeOS5yTQQnLYq7BOHpmjuC-kUejUU/edit?usp=sharing.

## Contributions

Contributed genes: IB, FKR, FM, SC, EW, PK

Analyzed data: LW, GZ

Wrote paper: LW, FKR, IB

Edited paper: FKR, FM, SC, EW, PK, GZ, LW, IB

## References

Abreu, Isidro, Ángela Saéz, Rosario Castro-Rodríguez, Viviana Escudero, Benjamín Rodríguez-Haas, Marta Senovilla, Camille Larue, et al. 2017. “Medicago Truncatula Zinc-Iron Permease6 Provides Zinc to Rhizobia-Infected Nodule Cells.” Plant, Cell & Environment 40 (11): 2706–19.

Aggarwal, Sipla, Anil Kumar, Kaushal K. Bhati, Gazaldeep Kaur, Vishnu Shukla, Siddharth Tiwari, and Ajay K. Pandey. 2018. “RNAi-Mediated Downregulation of Inositol Pentakisphosphate Kinase (IPK1) in Wheat Grains Decreases Phytic Acid Levels and Increases Fe and Zn Accumulation.” Frontiers in Plant Science 9 (March): 259.

Ahmad, Izhar, Jean Devonshire, Radwa Mohamed, Michael Schultze, and Frans J. M. Maathuis. 2016. “Overexpression of the Potassium Channel TPKb in Small Vacuoles Confers Osmotic and Drought Tolerance to Rice.” The New Phytologist 209 (3): 1040–48.

Ahmad, Izhar, Afaq Mian, and Frans J. M. Maathuis. 2016. “Overexpression of the Rice AKT1 Potassium Channel Affects Potassium Nutrition and Rice Drought Tolerance.” Journal of Experimental Botany 67 (9): 2689–98.

Andrés-Colás, Nuria, Vicente Sancenón, Susana Rodríguez-Navarro, Sonia Mayo, Dennis J. Thiele, Joseph R. Ecker, Sergi Puig, and Lola Peñarrubia. 2006. “The Arabidopsis Heavy Metal P-Type ATPase HMA5 Interacts with Metallochaperones and Functions in Copper Detoxification of Roots.” The Plant Journal: For Cell and Molecular Biology 45 (2): 225–36.

Arrivault, Stéphanie, Toralf Senger, and Ute Krämer. 2006. “The Arabidopsis Metal Tolerance Protein AtMTP3 Maintains Metal Homeostasis by Mediating Zn Exclusion from the Shoot under Fe Deficiency and Zn Oversupply.” The Plant Journal: For Cell and Molecular Biology 46 (5): 861–79.

Ashburner, M., C. A. Ball, J. A. Blake, D. Botstein, H. Butler, J. M. Cherry, A. P. Davis, et al. 2000. “Gene Ontology: Tool for the Unification of Biology. The Gene Ontology Consortium.” Nature Genetics 25 (1): 25–29.

Assunção, Ana G. L., Eva Herrero, Ya-Fen Lin, Bruno Huettel, Sangita Talukdar, Cezary Smaczniak, Richard G. H. Immink, et al. 2010. “Arabidopsis Thaliana Transcription Factors bZIP19 and bZIP23 Regulate the Adaptation to Zinc Deficiency.” Proceedings of the National Academy of Sciences of the United States of America 107 (22): 10296–301.

Barberon, Marie, Guillaume Dubeaux, Cornelia Kolb, Erika Isono, Enric Zelazny, and Grégory Vert. 2014. “Polarization of IRON-REGULATED TRANSPORTER 1 (IRT1) to the Plant-Soil Interface Plays Crucial Role in Metal Homeostasis.” Proceedings of the National Academy of Sciences of the United States of America 111 (22): 8293–98.

Barkla, Bronwyn J., and Omar Pantoja. 1996. “PHYSIOLOGY OF ION TRANSPORT ACROSS THE TONOPLAST OF HIGHER PLANTS.” Annual Review of Plant Physiology and Plant Molecular Biology 47 (June): 159–84.

Bashir, Khurram, Yasuhiro Ishimaru, Hugo Shimo, Seiji Nagasaka, Masaru Fujimoto, Hideki Takanashi, Nobuhiro Tsutsumi, Gynheung An, Hiromi Nakanishi, and Naoko K. Nishizawa. 2011. “The Rice Mitochondrial Iron Transporter Is Essential for Plant Growth.” Nature Communications 2: 322.

Baxter, I. R., O. Vitek, B. Lahner, B. Muthukumar, M. Borghi, J. Morrissey, M. L. Guerinot, and D. E. Salt. 2008. “The Leaf Ionome as a Multivariable System to Detect a Plant’s Physiological Status.” Proceedings of the National Academy of Sciences of the United States of America 105 (33): 12081–86.

Baxter, Ivan. 2020. “We Aren’t Good at Picking Candidate Genes, and It’s Slowing Us down.” Current Opinion in Plant Biology 54 (April): 57–60.

Baxter, Ivan, Jessica N. Brazelton, Danni Yu, Yu S. Huang, Brett Lahner, Elena Yakubova, Yan Li, et al. 2010. “A Coastal Cline in Sodium Accumulation in Arabidopsis Thaliana Is Driven by Natural Variation of the Sodium Transporter AtHKT1;1.” PLoS Genetics 6 (11): e1001193.

Baxter, Ivan, Prashant S. Hosmani, Ana Rus, Brett Lahner, Justin O. Borevitz, Balasubramaniam Muthukumar, Michael V. Mickelbart, Lukas Schreiber, Rochus B. Franke, and David E. Salt. 2009. “Root Suberin Forms an Extracellular Barrier That Affects Water Relations and Mineral Nutrition in Arabidopsis.” PLoS Genetics 5 (5): e1000492.

Baxter, Ivan, Balasubramaniam Muthukumar, Hyeong Cheol Park, Peter Buchner, Brett Lahner, John Danku, Keyan Zhao, et al. 2008. “Variation in Molybdenum Content across Broadly Distributed Populations of Arabidopsis Thaliana Is Controlled by a Mitochondrial Molybdenum Transporter (MOT1).” PLoS Genetics 4 (2): e1000004.

Bernal, María, David Casero, Vasantika Singh, Grandon T. Wilson, Arne Grande, Huijun Yang, Sheel C. Dodani, et al. 2012. “Transcriptome Sequencing Identifies SPL7-Regulated Copper Acquisition Genes FRO4/FRO5 and the Copper Dependence of Iron Homeostasis in Arabidopsis.” The Plant Cell 24 (2): 738–61.

Bhati, Kaushal Kumar, Anshu Alok, Anil Kumar, Jagdeep Kaur, Siddharth Tiwari, and Ajay Kumar Pandey. 2016. “Silencing of ABCC13 Transporter in Wheat Reveals Its Involvement in Grain Development, Phytic Acid Accumulation and Lateral Root Formation.” Journal of Experimental Botany 67 (14): 4379–89.

Borghi, Monica, Ana Rus, and David E. Salt. 2011. “Loss-of-Function of Constitutive Expresser of Pathogenesis Related Genes5 Affects Potassium Homeostasis in Arabidopsis Thaliana.” PloS One 6 (10): e26360.

Cailliatte, Rémy, Adam Schikora, Jean-François Briat, Stéphane Mari, and Catherine Curie. 2010. “High-Affinity Manganese Uptake by the Metal Transporter NRAMP1 Is Essential for Arabidopsis Growth in Low Manganese Conditions.” The Plant Cell 22 (3): 904–17.

Cao, Min-Jie, Zhen Wang, Markus Wirtz, Ruediger Hell, David J. Oliver, and Cheng-Bin Xiang. 2013. “SULTR3;1 Is a Chloroplast-Localized Sulfate Transporter in Arabidopsis Thaliana.” The Plant Journal: For Cell and Molecular Biology 73 (4): 607–16.

Carbon, Seth, Amelia Ireland, Christopher J. Mungall, Shengqiang Shu, Brad Marshall, Suzanna Lewis, AmiGO Hub, and Web Presence Working Group. 2009. “AmiGO: Online Access to Ontology and Annotation Data.” Bioinformatics 25 (2): 288–89.

Catarecha, Pablo, Maria Dolores Segura, José Manuel Franco-Zorrilla, Berenice García-Ponce, Mónica Lanza, Roberto Solano, Javier Paz-Ares, and Antonio Leyva. 2007. “A Mutant of the Arabidopsis Phosphate Transporter PHT1;1 Displays Enhanced Arsenic Accumulation.” The Plant Cell 19 (3): 1123–33.

Chan-Rodriguez, David, and Elsbeth L. Walker. 2018. “Analysis of Yellow Striped Mutants of Zea Mays Reveals Novel Loci Contributing to Iron Deficiency Chlorosis.” Frontiers in Plant Science 9 (February): 157.

Chao, Dai-Yin, Patrycja Baraniecka, John Danku, Anna Koprivova, Brett Lahner, Hongbing Luo, Elena Yakubova, Brian Dilkes, Stanislav Kopriva, and David E. Salt. 2014. “Variation in Sulfur and Selenium Accumulation Is Controlled by Naturally Occurring Isoforms of the Key Sulfur Assimilation Enzyme ADENOSINE 5’-PHOSPHOSULFATE REDUCTASE2 across the Arabidopsis Species Range.” Plant Physiology 166 (3): 1593–1608.

Chao, Dai-Yin, Yi Chen, Jiugeng Chen, Shulin Shi, Ziru Chen, Chengcheng Wang, John M. Danku, Fang-Jie Zhao, and David E. Salt. 2014. “Genome-Wide Association Mapping Identifies a New Arsenate Reductase Enzyme Critical for Limiting Arsenic Accumulation in Plants.” PLoS Biology 12 (12): e1002009.

Chao, Dai-Yin, Kenneth Gable, Ming Chen, Ivan Baxter, Charles R. Dietrich, Edgar B. Cahoon, Mary Lou Guerinot, et al. 2011. “Sphingolipids in the Root Play an Important Role in Regulating the Leaf Ionome in Arabidopsis Thaliana.” The Plant Cell 23 (3): 1061–81.

Chao, Dai-Yin, Adriano Silva, Ivan Baxter, Yu S. Huang, Magnus Nordborg, John Danku, Brett Lahner, Elena Yakubova, and David E. Salt. 2012. “Genome-Wide Association Studies Identify Heavy Metal ATPase3 as the Primary Determinant of Natural Variation in Leaf Cadmium in Arabidopsis Thaliana.” PLoS Genetics 8 (9): e1002923.

Che, Jing, Kengo Yokosho, Naoki Yamaji, and Jian Feng Ma. 2019. “A Vacuolar Phytosiderophore Transporter Alters Iron and Zinc Accumulation in Polished Rice Grains.” Plant Physiology 181 (1): 276–88.

Chen, Jieyu, Yu Liu, Jun Ni, Yifeng Wang, Youhuang Bai, Jing Shi, Jian Gan, Zhongchang Wu, and Ping Wu. 2011. “OsPHF1 Regulates the Plasma Membrane Localization of Low-and High-Affinity Inorganic Phosphate Transporters and Determines Inorganic Phosphate Uptake and Translocation in Rice.” Plant Physiology 157 (1): 269–78.

Chen, Yi, Sheng-Kai Sun, Zhong Tang, Guidong Liu, Katie L. Moore, Frans J. M. Maathuis, Antony J. Miller, Steve P. McGrath, and Fang-Jie Zhao. 2017. “The Nodulin 26-like Intrinsic Membrane Protein OsNIP3;2 Is Involved in Arsenite Uptake by Lateral Roots in Rice.” Journal of Experimental Botany 68 (11): 3007–16.

Chen, Zhi Chang, Naoki Yamaji, Tomoaki Horie, Jing Che, Jian Li, Gynheung An, and Jian Feng Ma. 2017. “A Magnesium Transporter OsMGT1 Plays a Critical Role in Salt Tolerance in Rice.” Plant Physiology 174 (3): 1837–49.

Chen, Zonghui, Yumi Fujii, Naoki Yamaji, Sakine Masuda, Yuma Takemoto, Takehiro Kamiya, Yusufujiang Yusuyin, et al. 2013. “Mn Tolerance in Rice Is Mediated by MTP8.1, a Member of the Cation Diffusion Facilitator Family.” Journal of Experimental Botany 64 (14): 4375–87.

Connorton, James M., Rachel E. Webster, Ninghui Cheng, and Jon K. Pittman. 2012. “Knockout of Multiple Arabidopsis cation/H(+) Exchangers Suggests Isoform-Specific Roles in Metal Stress Response, Germination and Seed Mineral Nutrition.” PloS One 7 (10): e47455.

Delhaize, E. 1996. “A Metal-Accumulator Mutant of Arabidopsis Thaliana.” Plant Physiology 111 (3): 849–55.

Deng, Fenglin, Naoki Yamaji, Jixing Xia, and Jian Feng Ma. 2013. “A Member of the Heavy Metal P-Type ATPase OsHMA5 Is Involved in Xylem Loading of Copper in Rice.” Plant Physiology 163 (3): 1353–62.

Desbrosses-Fonrouge, Anne-Garlonn, Katrin Voigt, Astrid Schröder, Stéphanie Arrivault, Sébastien Thomine, and Ute Krämer. 2005. “Arabidopsis Thaliana MTP1 Is a Zn Transporter in the Vacuolar Membrane Which Mediates Zn Detoxification and Drives Leaf Zn Accumulation.” FEBS Letters 579 (19): 4165–74.

Devaiah, Ballachanda N., Ramaiah Madhuvanthi, Athikkattuvalasu S. Karthikeyan, and Kashchandra G. Raghothama. 2009. “Phosphate Starvation Responses and Gibberellic Acid Biosynthesis Are Regulated by the MYB62 Transcription Factor in Arabidopsis.” Molecular Plant 2 (1): 43–58.

Dhankher, Om Parkash, Barry P. Rosen, Elizabeth C. McKinney, and Richard B. Meagher. 2006. “Hyperaccumulation of Arsenic in the Shoots of Arabidopsis Silenced for Arsenate Reductase (ACR2).” Proceedings of the National Academy of Sciences of the United States of America 103 (14): 5413–18.

Durbak, Amanda R., Kimberly A. Phillips, Sharon Pike, Malcolm A. O’Neill, Jonathan Mares, Andrea Gallavotti, Simon T. Malcomber, Walter Gassmann, and Paula McSteen. 2014. “Transport of Boron by the Tassel-less1 Aquaporin Is Critical for Vegetative and Reproductive Development in Maize.” The Plant Cell 26 (7): 2978–95.

Eide, D., M. Broderius, J. Fett, and M. L. Guerinot. 1996. “A Novel Iron-Regulated Metal Transporter from Plants Identified by Functional Expression in Yeast.” Proceedings of the National Academy of Sciences of the United States of America 93 (11): 5624–28.

Eroglu, Seckin, Ricardo F. H. Giehl, Bastian Meier, Michiko Takahashi, Yasuko Terada, Konstantin Ignatyev, Elisa Andresen, Hendrik Küpper, Edgar Peiter, and Nicolaus von Wirén. 2017. “Metal Tolerance Protein 8 Mediates Manganese Homeostasis and Iron Reallocation during Seed Development and Germination.” Plant Physiology 174 (3): 1633–47.

Eroglu, Seckin, Bastian Meier, Nicolaus von Wirén, and Edgar Peiter. 2016. “The Vacuolar Manganese Transporter MTP8 Determines Tolerance to Iron Deficiency-Induced Chlorosis in Arabidopsis.” Plant Physiology 170 (2): 1030–45.

Gao, Yi-Qun, Jiu-Geng Chen, Zi-Ru Chen, Dong An, Qiao-Yan Lv, Mei-Ling Han, Ya-Ling Wang, David E. Salt, and Dai-Yin Chao. 2017. “A New Vesicle Trafficking Regulator CTL1 Plays a Crucial Role in Ion Homeostasis.” PLoS Biology 15 (12): e2002978.

Gil-Díez, Patricia, Manuel Tejada-Jiménez, Javier León-Mediavilla, Jiangqi Wen, Kirankumar S. Mysore, Juan Imperial, and Manuel González-Guerrero. 2018. “MtMOT1.2 Is Responsible for Molybdate Supply to Medicago Truncatula Nodules.” Plant, Cell & Environment, June. https://doi.org/10.1111/pce.13388.

Gobert, Anthony, Graeme Park, Anna Amtmann, Dale Sanders, and Frans J. M. Maathuis. 2006. “Arabidopsis Thaliana Cyclic Nucleotide Gated Channel 3 Forms a Non-Selective Ion Transporter Involved in Germination and Cation Transport.” Journal of Experimental Botany 57 (4): 791–800.

Guo, Jiangbo, Xiaojing Dai, Wenzhong Xu, and Mi Ma. 2008. “Overexpressing GSH1 and AsPCS1 Simultaneously Increases the Tolerance and Accumulation of Cadmium and Arsenic in Arabidopsis Thaliana.” Chemosphere 72 (7): 1020–26.

Guo, Kun Mei, Olga Babourina, David A. Christopher, Tamas Borsic, and Zed Rengel. 2010. “The Cyclic Nucleotide-Gated Channel AtCNGC10 Transports Ca2+ and Mg2+ in Arabidopsis.” Physiologia Plantarum 139 (3): 303–12.

Harris, M. A., J. Clark, A. Ireland, J. Lomax, M. Ashburner, R. Foulger, K. Eilbeck, et al. 2004. “The Gene Ontology (GO) Database and Informatics Resource.” Nucleic Acids Research 32 (Database issue): D258–61.

Hayashi, Shimpei, Masato Kuramata, Tadashi Abe, Hiroki Takagi, Kenjirou Ozawa, and Satoru Ishikawa. 2017. “Phytochelatin Synthase OsPCS1 Plays a Crucial Role in Reducing Arsenic Levels in Rice Grains.” The Plant Journal: For Cell and Molecular Biology 91 (5): 840–48.

Haydon, Michael J., Miki Kawachi, Markus Wirtz, Stefan Hillmer, Rüdiger Hell, and Ute Krämer. 2012. “Vacuolar Nicotianamine Has Critical and Distinct Roles under Iron Deficiency and for Zinc Sequestration in Arabidopsis.” The Plant Cell 24 (2): 724–37.

Hindt, Maria N., Garo Z. Akmakjian, Kara L. Pivarski, Tracy Punshon, Ivan Baxter, David E. Salt, and Mary Lou Guerinot. 2017. “BRUTUS and Its Paralogs, BTS LIKE1 and BTS LIKE2, Encode Important Negative Regulators of the Iron Deficiency Response in Arabidopsis Thaliana.” Metallomics: Integrated Biometal Science 9 (7): 876–90.

Horie, Tomoaki, Alex Costa, Tae Houn Kim, Min Jung Han, Rie Horie, Ho-Yin Leung, Akio Miyao, Hirohiko Hirochika, Gynheung An, and Julian I. Schroeder. 2007. “Rice OsHKT2;1 Transporter Mediates Large Na+ Influx Component into K+-Starved Roots for Growth.” The EMBO Journal 26 (12): 3003–14.

Huang, Xin-Yuan, Dai-Yin Chao, Anna Koprivova, John Danku, Markus Wirtz, Steffen Müller, Francisco J. Sandoval, et al. 2016. “Nuclear Localised MORE SULPHUR ACCUMULATION1 Epigenetically Regulates Sulphur Homeostasis in Arabidopsis Thaliana.” PLoS Genetics 12 (9): e1006298.

Huang, Xin-Yuan, Fenglin Deng, Naoki Yamaji, Shannon R. M. Pinson, Miho Fujii-Kashino, John Danku, Alex Douglas, Mary Lou Guerinot, David E. Salt, and Jian Feng Ma. 2016. “A Heavy Metal P-Type ATPase OsHMA4 Prevents Copper Accumulation in Rice Grain.” Nature Communications 7 (July): 12138.

Huang, Xin-Yuan, Huan Liu, Yu-Fei Zhu, Shannon R. M. Pinson, Hong-Xuan Lin, Mary Lou Guerinot, Fang-Jie Zhao, and David E. Salt. 2019. “Natural Variation in a Molybdate Transporter Controls Grain Molybdenum Concentration in Rice.” The New Phytologist 221 (4): 1983–97.

Hussain, Dawar, Michael J. Haydon, Yuwen Wang, Edwin Wong, Sarah M. Sherson, Jeff Young, James Camakaris, Jeffrey F. Harper, and Christopher S. Cobbett. 2004. “P-Type ATPase Heavy Metal Transporters with Roles in Essential Zinc Homeostasis in Arabidopsis.” The Plant Cell 16 (5): 1327–39.

Isayenkov, Stanislav V., and Frans J. M. Maathuis. 2008. “The Arabidopsis Thaliana Aquaglyceroporin AtNIP7;1 Is a Pathway for Arsenite Uptake.” FEBS Letters 582 (11): 1625–28.

Ishimaru, Yasuhiro, Khurram Bashir, Masaru Fujimoto, Gynheung An, Reiko Nakanishi Itai, Nobuhiro Tsutsumi, Hiromi Nakanishi, and Naoko K. Nishizawa. 2009. “Rice-Specific Mitochondrial Iron-Regulated Gene (MIR) Plays an Important Role in Iron Homeostasis.” Molecular Plant 2 (5): 1059–66.

Ishimaru, Yasuhiro, Hiroshi Masuda, Khurram Bashir, Haruhiko Inoue, Takashi Tsukamoto, Michiko Takahashi, Hiromi Nakanishi, et al. 2010. “Rice Metal-Nicotianamine Transporter, OsYSL2, Is Required for the Long-Distance Transport of Iron and Manganese.” The Plant Journal: For Cell and Molecular Biology 62 (3): 379–90.

Kamiya, Takehiro, Monica Borghi, Peng Wang, John M. C. Danku, Lothar Kalmbach, Prashant S. Hosmani, Sadaf Naseer, Toru Fujiwara, Niko Geldner, and David E. Salt. 2015. “The MYB36 Transcription Factor Orchestrates Casparian Strip Formation.” Proceedings of the National Academy of Sciences of the United States of America 112 (33): 10533–38.

Kamiya, Takehiro, Mutsumi Yamagami, Masami Yokota Hirai, and Toru Fujiwara. 2012. “Establishment of an in Planta Magnesium Monitoring System Using CAX3 Promoter-Luciferase in Arabidopsis.” Journal of Experimental Botany 63 (1): 355–63.

Khan, Ghazanfar Abbas, Samir Bouraine, Stefanie Wege, Yuanyuan Li, Matthieu de Carbonnel, Pierre Berthomieu, Yves Poirier, and Hatem Rouached. 2014. “Coordination between Zinc and Phosphate Homeostasis Involves the Transcription Factor PHR1, the Phosphate Exporter PHO1, and Its Homologue PHO1;H3 in Arabidopsis.” Journal of Experimental Botany 65 (3): 871–84.

Kim, Do-Young, Lucien Bovet, Masayoshi Maeshima, Enrico Martinoia, and Youngsook Lee. 2007. “The ABC Transporter AtPDR8 Is a Cadmium Extrusion Pump Conferring Heavy Metal Resistance: Role of AtPDR8 in Cadmium Resistance.” The Plant Journal: For Cell and Molecular Biology 50 (2): 207–18.

Kim, Sun A., Tracy Punshon, Antonio Lanzirotti, Liangtao Li, José M. Alonso, Joseph R. Ecker, Jerry Kaplan, and Mary Lou Guerinot. 2006. “Localization of Iron in Arabidopsis Seed Requires the Vacuolar Membrane Transporter VIT1 Science 314 (5803): 1295–98.

Kim, Yu-Young, Hyunju Choi, Shoji Segami, Hyung-Taeg Cho, Enrico Martinoia, Masayoshi Maeshima, and Youngsook Lee. 2009. “AtHMA1 Contributes to the Detoxification of Excess Zn(II) in Arabidopsis.” The Plant Journal: For Cell and Molecular Biology 58 (5): 737–53.

Kisko, Mushtak, Nadia Bouain, Alaeddine Safi, Anna Medici, Robert C. Akkers, David Secco, Gilles Fouret, et al. 2018. “LPCAT1 Controls Phosphate Homeostasis in a Zinc-Dependent Manner.” eLife 7 (February). https://doi.org/10.7554/eLife.32077.

Klaumann, Sandra, Sebastian D. Nickolaus, Sarah H. Fürst, Sabrina Starck, Sabine Schneider, H. Ekkehard Neuhaus, and Oliver Trentmann. 2011. “The Tonoplast Copper Transporter COPT5 Acts as an Exporter and Is Required for Interorgan Allocation of Copper in Arabidopsis Thaliana.” The New Phytologist 192 (2): 393–404.

Kobayashi, Natsuko I., Naoki Yamaji, Hiroki Yamamoto, Kaoru Okubo, Hiroki Ueno, Alex Costa, Keitaro Tanoi, et al. 2017. “OsHKT1;5 Mediates Na+ Exclusion in the Vasculature to Protect Leaf Blades and Reproductive Tissues from Salt Toxicity in Rice.” The Plant Journal: For Cell and Molecular Biology 91 (4): 657–70.

Kobayashi, Takanori, Seiji Nagasaka, Takeshi Senoura, Reiko Nakanishi Itai, Hiromi Nakanishi, and Naoko K. Nishizawa. 2013. “Iron-Binding Haemerythrin RING Ubiquitin Ligases Regulate Plant Iron Responses and Accumulation.” Nature Communications 4: 2792.

Koprivova, Anna, Marco Giovannetti, Patrycja Baraniecka, Bok-Rye Lee, Cécile Grondin, Olivier Loudet, and Stanislav Kopriva. 2013. “Natural Variation in the ATPS1 Isoform of ATP Sulfurylase Contributes to the Control of Sulfate Levels in Arabidopsis.” Plant Physiology 163 (3): 1133–41.

Kühnlenz, Tanja, Christian Hofmann, Shimpei Uraguchi, Holger Schmidt, Stefanie Schempp, Michael Weber, Brett Lahner, David E. Salt, and Stephan Clemens. 2016. “Phytochelatin Synthesis Promotes Leaf Zn Accumulation of Arabidopsis Thaliana Plants Grown in Soil with Adequate Zn Supply and Is Essential for Survival on Zn-Contaminated Soil.” Plant & Cell Physiology 57 (11): 2342–52.

Lagarde, Delphine, Mireille Basset, Marc Lepetit, Geneviève Conejero, Frédéric Gaymard, Suzette Astruc, and Claude Grignon. 1996. “Tissue-Specific Expression of Arabidopsis AKT1 Gene Is Consistent with a Role in K+ Nutrition.” The Plant Journal: For Cell and Molecular Biology 9 (2): 195–203.

Lanquar, Viviane, Magali Schnell Ramos, Françoise Lelièvre, Hélène Barbier-Brygoo, Anja Krieger-Liszkay, Ute Krämer, and Sébastien Thomine. 2010. “Export of Vacuolar Manganese by AtNRAMP3 and AtNRAMP4 Is Required for Optimal Photosynthesis and Growth under Manganese Deficiency.” Plant Physiology 152 (4): 1986–99.

Laurie, Sophie, Kevin A. Feeney, Frans J. M. Maathuis, Peter J. Heard, Sherralyn J. Brown, and Roger A. Leigh. 2002. “A Role for HKT1 in Sodium Uptake by Wheat Roots.” The Plant Journal: For Cell and Molecular Biology 32 (2): 139–49.

Lee, Sichul, Jeff C. Chiecko, Sun A. Kim, Elsbeth L. Walker, Youngsook Lee, Mary Lou Guerinot, and Gynheung An. 2009. “Disruption of OsYSL15 Leads to Iron Inefficiency in Rice Plants.” Plant Physiology 150 (2): 786–800.

Lee, Sichul, Hee Joong Jeong, Sun A. Kim, Joohyun Lee, Mary Lou Guerinot, and Gynheung An. 2010. “OsZIP5 Is a Plasma Membrane Zinc Transporter in Rice.” Plant Molecular Biology 73 (4-5): 507–17.

Lee, Sichul, You-Sun Kim, Un Sil Jeon, Yoon-Keun Kim, Jan K. Schjoerring, and Gynheung An. 2012. “Activation of Rice Nicotianamine Synthase 2 (OsNAS2) Enhances Iron Availability for Biofortification.” Molecules and Cells 33 (3): 269–75.

León-Mediavilla, Javier, Marta Senovilla, Jesús Montiel, Patricia Gil-Díez, Ángela Saez, Igor S. Kryvoruchko, María Reguera, Michael K. Udvardi, Juan Imperial, and Manuel González-Guerrero. 2018. “MtMTP2-Facilitated Zinc Transport Into Intracellular Compartments Is Essential for Nodule Development in Medicago Truncatula.” Frontiers in Plant Science 9 (July): 990.

Li, Hong, Miao Yu, Xin-Qiao Du, Zhi-Fang Wang, Wei-Hua Wu, Francisco J. Quintero, Xue-Hua Jin, Hao-Dong Li, and Yi Wang. 2017. “NRT1.5/NPF7.3 Functions as a Proton-Coupled H+/K+ Antiporter for K+ Loading into the Xylem in Arabidopsis.” The Plant Cell 29 (8): 2016–26.

Lindsay, Emma R., and Frans J. M. Maathuis. 2016. “Arabidopsis Thaliana NIP7;1 Is Involved in Tissue Arsenic Distribution and Tolerance in Response to Arsenate.” FEBS Letters 590 (6): 779–86.

Lin, Ya-Fen, Hong-Ming Liang, Shu-Yi Yang, Annegret Boch, Stephan Clemens, Chyi-Chuann Chen, Jing-Fen Wu, Jing-Ling Huang, and Kuo-Chen Yeh. 2009. “Arabidopsis IRT3 Is a Zinc-Regulated and Plasma Membrane Localized Zinc/iron Transporter.” The New Phytologist 182 (2): 392–404.

Liu, Tzu-Yin, Teng-Kuei Huang, Ching-Ying Tseng, Ya-Shiuan Lai, Shu-I Lin, Wei-Yi Lin, June-Wei Chen, and Tzyy-Jen Chiou. 2012. “PHO2-Dependent Degradation of PHO1 Modulates Phosphate Homeostasis in Arabidopsis.” The Plant Cell 24 (5): 2168–83.

Liu, Xiao-Min, Jonguk An, Hay Ju Han, Sun Ho Kim, Chae Oh Lim, Dae-Jin Yun, and Woo Sik Chung. 2014. “ZAT11, a Zinc Finger Transcription Factor, Is a Negative Regulator of Nickel Ion Tolerance in Arabidopsis.” Plant Cell Reports 33 (12): 2015–21.

Li, Xiaoli, Huimin Zhang, Qin Ai, Gang Liang, and Diqiu Yu. 2016. “Two bHLH Transcription Factors, bHLH34 and bHLH104, Regulate Iron Homeostasis in Arabidopsis Thaliana.” Plant Physiology 170 (4): 2478–93.

Long, Terri A., Hironaka Tsukagoshi, Wolfgang Busch, Brett Lahner, David E. Salt, and Philip N. Benfey. 2010. “The bHLH Transcription Factor POPEYE Regulates Response to Iron Deficiency in Arabidopsis Roots.” The Plant Cell 22 (7): 2219–36.

Loudet, Olivier, Vera Saliba-Colombani, Christine Camilleri, Fanny Calenge, Virginie Gaudon, Anna Koprivova, Kathryn A. North, Stanislav Kopriva, and Françoise Daniel-Vedele. 2007. “Natural Variation for Sulfate Content in Arabidopsis Thaliana Is Highly Controlled by APR2.” Nature Genetics 39 (7): 896–900.

Mao, Dandan, Jian Chen, Lianfu Tian, Zhenhua Liu, Lei Yang, Renjie Tang, Jian Li, et al. 2014. “Arabidopsis Transporter MGT6 Mediates Magnesium Uptake and Is Required for Growth under Magnesium Limitation.” The Plant Cell 26 (5): 2234–48.

McDowell, Stephen C., Garo Akmakjian, Chris Sladek, David Mendoza-Cozatl, Joe B. Morrissey, Nick Saini, Ron Mittler, et al. 2013. “Elemental Concentrations in the Seed of Mutants and Natural Variants of Arabidopsis Thaliana Grown under Varying Soil Conditions.” PloS One 8 (5): e63014.

Mi, Huaiyu, Xiaosong Huang, Anushya Muruganujan, Haiming Tang, Caitlin Mills, Diane Kang, and Paul D. Thomas. 2017. “PANTHER Version 11: Expanded Annotation Data from Gene Ontology and Reactome Pathways, and Data Analysis Tool Enhancements.” Nucleic Acids Research 45 (D1): D183–89.

Milner, Matthew J., Jesse Seamon, Eric Craft, and Leon V. Kochian. 2013. “Transport Properties of Members of the ZIP Family in Plants and Their Role in Zn and Mn Homeostasis.” Journal of Experimental Botany 64 (1): 369–81.

Miwa, Kyoko, Junpei Takano, and Toru Fujiwara. 2006. “Improvement of Seed Yields under Boron-Limiting Conditions through Overexpression of BOR1, a Boron Transporter for Xylem Loading, in Arabidopsis Thaliana.” The Plant Journal: For Cell and Molecular Biology 46 (6): 1084–91.

Miwa, Kyoko, Shinji Wakuta, Shigeki Takada, Koji Ide, Junpei Takano, Satoshi Naito, Hiroyuki Omori, Toshiro Matsunaga, and Toru Fujiwara. 2013. “Roles of BOR2, a Boron Exporter, in Cross Linking of Rhamnogalacturonan II and Root Elongation under Boron Limitation in Arabidopsis.” Plant Physiology 163 (4): 1699–1709.

Morrissey, Joe, Ivan R. Baxter, Joohyun Lee, Liangtao Li, Brett Lahner, Natasha Grotz, Jerry Kaplan, David E. Salt, and Mary Lou Guerinot. 2009. “The Ferroportin Metal Efflux Proteins Function in Iron and Cobalt Homeostasis in Arabidopsis.” The Plant Cell 21 (10): 3326–38.

Nagarajan, Vinay K., Ajay Jain, Michael D. Poling, Anthony J. Lewis, Kashchandra G. Raghothama, and Aaron P. Smith. 2011. “Arabidopsis Pht1;5 Mobilizes Phosphate between Source and Sink Organs and Influences the Interaction between Phosphate Homeostasis and Ethylene Signaling.” Plant Physiology 156 (3): 1149–63.

Nakagawa, Yuko, Hideki Hanaoka, Masaharu Kobayashi, Kazumaru Miyoshi, Kyoko Miwa, and Toru Fujiwara. 2007. “Cell-Type Specificity of the Expression of Os BOR1, a Rice Efflux Boron Transporter Gene, Is Regulated in Response to Boron Availability for Efficient Boron Uptake and Xylem Loading.” The Plant Cell 19 (8): 2624–35.

Nilsson, Lena, Renate Müller, and Tom Hamborg Nielsen. 2007. “Increased Expression of the MYB-Related Transcription Factor, PHR1, Leads to Enhanced Phosphate Uptake in Arabidopsis Thaliana.” Plant, Cell & Environment 30 (12): 1499–1512.

Olsen, Lene Irene, Thomas H. Hansen, Camille Larue, Jeppe Thulin Østerberg, Robert D. Hoffmann, Johannes Liesche, Ute Krämer, et al. 2016. “Mother-Plant-Mediated Pumping of Zinc into the Developing Seed.” Nature Plants 2 (5): 16036.

Palmer, Christine M., Maria N. Hindt, Holger Schmidt, Stephan Clemens, and Mary Lou Guerinot. 2013. “MYB10 and MYB72 Are Required for Growth under Iron-Limiting Conditions.” PLoS Genetics 9 (11): e1003953.

Park, Jiyoung, Won-Yong Song, Donghwi Ko, Yujin Eom, Thomas H. Hansen, Michaela Schiller, Tai Gyu Lee, Enrico Martinoia, and Youngsook Lee. 2012. “The Phytochelatin Transporters AtABCC1 and AtABCC2 Mediate Tolerance to Cadmium and Mercury: ABC Transporters for PC-Dependent Cd and Hg Tolerance.” The Plant Journal: For Cell and Molecular Biology 69 (2): 278–88.

Peiter, Edgar, Barbara Montanini, Anthony Gobert, Pai Pedas, Søren Husted, Frans J. M. Maathuis, Damien Blaudez, Michel Chalot, and Dale Sanders. 2007. “A Secretory Pathway-Localized Cation Diffusion Facilitator Confers Plant Manganese Tolerance.” Proceedings of the National Academy of Sciences of the United States of America 104 (20): 8532–37.

Pita-Barbosa, Alice, Felipe K. Ricachenevsky, Michael Wilson, Tania Dottorini, and David E. Salt. 2019. “Transcriptional Plasticity Buffers Genetic Variation in Zinc Homeostasis.” Scientific Reports 9 (1): 19482.

Pottier, Mathieu, Jean Dumont, Céline Masclaux-Daubresse, and Sébastien Thomine. 2019. “Autophagy Is Essential for Optimal Translocation of Iron to Seeds in Arabidopsis.” Journal of Experimental Botany 70 (3): 859–69.

Qin, Ya-Juan, Wei-Hua Wu, and Yi Wang. 2019. “ZmHAK5 and ZmHAK1 Function in K+ Uptake and Distribution in Maize under Low K+ Conditions.” Journal of Integrative Plant Biology 61 (6): 691–705.

Qi, Zhi, Corrina R. Hampton, Ryoung Shin, Bronwyn J. Barkla, Philip J. White, and Daniel P. Schachtman. 2008. “The High Affinity K+ Transporter AtHAK5 Plays a Physiological Role in Planta at Very Low K+ Concentrations and Provides a Caesium Uptake Pathway in Arabidopsis.” Journal of Experimental Botany 59 (3): 595–607.

Rai, Hiroki, Saki Yokoyama, Namiko Satoh-Nagasawa, Jun Furukawa, Takiko Nomi, Yasuka Ito, Shigeto Fujimura, et al. 2017. “Cesium Uptake by Rice Roots Largely Depends Upon a Single Gene, HAK1, Which Encodes a Potassium Transporter.” Plant & Cell Physiology 58 (9): 1486–93.

Rampey, Rebekah A., Andrew W. Woodward, Brianne N. Hobbs, Megan P. Tierney, Brett Lahner, David E. Salt, and Bonnie Bartel. 2006. “An Arabidopsis Basic Helix-Loop-Helix Leucine Zipper Protein Modulates Metal Homeostasis and Auxin Conjugate Responsiveness.” Genetics 174 (4): 1841–57.

Remm, M., C. E. Storm, and E. L. Sonnhammer. 2001. “Automatic Clustering of Orthologs and in-Paralogs from Pairwise Species Comparisons.” Journal of Molecular Biology 314 (5): 1041–52.

Remy, E., T. R. Cabrito, R. A. Batista, M. C. Teixeira, I. Sá-Correia, and P. Duque. 2012. “The Pht1;9 and Pht1;8 Transporters Mediate Inorganic Phosphate Acquisition by the Arabidopsis Thaliana Root during Phosphorus Starvation.” The New Phytologist 195 (2): 356–71.

Remy, Estelle, Tânia R. Cabrito, Rita A. Batista, Miguel C. Teixeira, Isabel Sá-Correia, and Paula Duque. 2015. “The Major Facilitator Superfamily Transporter ZIFL2 Modulates Cesium and Potassium Homeostasis in Arabidopsis.” Plant & Cell Physiology 56 (1): 148–62.

Ren, Xiao-Ling, Guo-Ning Qi, Han-Qian Feng, Shuai Zhao, Shuang-Shuang Zhao, Yi Wang, and Wei-Hua Wu. 2013. “Calcineurin B-like Protein CBL10 Directly Interacts with AKT1 and Modulates K+ Homeostasis in Arabidopsis.” The Plant Journal: For Cell and Molecular Biology 74 (2): 258–66.

Ren, Yujun, Yang Liu, Hongyu Chen, Gang Li, Xuelian Zhang, and Jie Zhao. 2012. “Type 4 Metallothionein Genes Are Involved in Regulating Zn Ion Accumulation in Late Embryo and in Controlling Early Seedling Growth in Arabidopsis.” Plant, Cell & Environment 35 (4): 770–89.

Ren, Zhong-Hai, Ji-Ping Gao, Le-Gong Li, Xiu-Ling Cai, Wei Huang, Dai-Yin Chao, Mei-Zhen Zhu, Zong-Yang Wang, Sheng Luan, and Hong-Xuan Lin. 2005. “A Rice Quantitative Trait Locus for Salt Tolerance Encodes a Sodium Transporter.” Nature Genetics 37 (10): 1141–46.

Robinson, Nigel J., Catherine M. Procter, Erin L. Connolly, and Mary Lou Guerinot. 1999. “A Ferric-Chelate Reductase for Iron Uptake from Soils.” Nature 397 (February): 694.

Rubio, Francisco, Manuel Nieves-Cordones, Fernando Alemán, and Vicente Martínez. 2008. “Relative Contribution of AtHAK5 and AtAKT1 to K+ Uptake in the High-Affinity Range of Concentrations.” Physiologia Plantarum 134 (4): 598–608.

Sancenón, Vicente, Sergi Puig, Isabel Mateu-Andrés, Eavan Dorcey, Dennis J. Thiele, and Lola Peñarrubia. 2004. “The Arabidopsis Copper Transporter COPT1 Functions in Root Elongation and Pollen Development.” The Journal of Biological Chemistry 279 (15): 15348–55.

Sanjaya, Pao-Yuan Hsiao, Ruey-Chih Su, Swee-Suak Ko, Chii-Gong Tong, Ray-Yu Yang, and Ming-Tsair Chan. 2008. “Overexpression of Arabidopsis Thaliana Tryptophan Synthase Beta 1 (AtTSB1) in Arabidopsis and Tomato Confers Tolerance to Cadmium Stress.” Plant, Cell & Environment 31 (8): 1074–85.

Sasaki, Akimasa, Naoki Yamaji, Namiki Mitani-Ueno, Miho Kashino, and Jian Feng Ma. 2015. “A Node-Localized Transporter OsZIP3 Is Responsible for the Preferential Distribution of Zn to Developing Tissues in Rice.” The Plant Journal: For Cell and Molecular Biology 84 (2): 374–84.

Sasaki, Akimasa, Naoki Yamaji, Kengo Yokosho, and Jian Feng Ma. 2012. “Nramp5 Is a Major Transporter Responsible for Manganese and Cadmium Uptake in Rice.” The Plant Cell 24 (5): 2155–67.

Schaaf, Gabriel, Annegret Honsbein, Anderson R. Meda, Silvia Kirchner, Daniel Wipf, and Nicolaus von Wirén. 2006. “AtIREG2 Encodes a Tonoplast Transport Protein Involved in Iron-Dependent Nickel Detoxification in Arabidopsis Thaliana Roots.” The Journal of Biological Chemistry 281 (35): 25532–40.

Secco, David, Arnaud Baumann, and Yves Poirier. 2010. “Characterization of the Rice PHO1 Gene Family Reveals a Key Role for OsPHO1;2 in Phosphate Homeostasis and the Evolution of a Distinct Clade in Dicotyledons.” Plant Physiology 152 (3): 1693–1704.

Selote, Devarshi, Rozalynne Samira, Anna Matthiadis, Jeffrey W. Gillikin, and Terri A. Long. 2015. “Iron-Binding E3 Ligase Mediates Iron Response in Plants by Targeting Basic Helix-Loop-Helix Transcription Factors.” Plant Physiology 167 (1): 273–86.

Senoura, Takeshi, Emi Sakashita, Takanori Kobayashi, Michiko Takahashi, May Sann Aung, Hiroshi Masuda, Hiromi Nakanishi, and Naoko K. Nishizawa. 2017. “The Iron-Chelate Transporter OsYSL9 Plays a Role in Iron Distribution in Developing Rice Grains.” Plant Molecular Biology 95 (4-5): 375–87.

Senovilla, Marta, Rosario Castro-Rodríguez, Isidro Abreu, Viviana Escudero, Igor Kryvoruchko, Michael K. Udvardi, Juan Imperial, and Manuel González-Guerrero. 2018. “Medicago Truncatula Copper Transporter 1 (MtCOPT1) Delivers Copper for Symbiotic Nitrogen Fixation.” The New Phytologist 218 (2): 696–709.

Shi, Huazhong, Byeong-Ha Lee, Shaw-Jye Wu, and Jian-Kang Zhu. 2003. “Overexpression of a Plasma Membrane Na+/H+ Antiporter Gene Improves Salt Tolerance in Arabidopsis Thaliana.” Nature Biotechnology 21 (1): 81–85.

Shin, Heungsop, Hwa-Soo Shin, Gary R. Dewbre, and Maria J. Harrison. 2004. “Phosphate Transport in Arabidopsis: Pht1;1 and Pht1;4 Play a Major Role in Phosphate Acquisition from Both Low-and High-Phosphate Environments.” The Plant Journal: For Cell and Molecular Biology 39 (4): 629–42.

Shin, Lung-Jiun, Jing-Chi Lo, and Kuo-Chen Yeh. 2012. “Copper Chaperone Antioxidant protein1 Is Essential for Copper Homeostasis.” Plant Physiology 159 (3): 1099–1110.

Song, Won-Yong, Kwan Sam Choi, Do Young Kim, Markus Geisler, Jiyoung Park, Vincent Vincenzetti, Maja Schellenberg, et al. 2010. “Arabidopsis PCR2 Is a Zinc Exporter Involved in Both Zinc Extrusion and Long-Distance Zinc Transport.” The Plant Cell 22 (7): 2237–52.

Song, Won-Yong, Tomohiro Yamaki, Naoki Yamaji, Donghwi Ko, Ki-Hong Jung, Miho Fujii-Kashino, Gynheung An, Enrico Martinoia, Youngsook Lee, and Jian Feng Ma. 2014. “A Rice ABC Transporter, OsABCC1, Reduces Arsenic Accumulation in the Grain.” Proceedings of the National Academy of Sciences of the United States of America 111 (44): 15699–704.

Stoeger, Thomas, Martin Gerlach, Richard I. Morimoto, and Luís A. Nunes Amaral. 2018. “Large-Scale Investigation of the Reasons Why Potentially Important Genes Are Ignored.” PLoS Biology 16 (9): e2006643.

Sunkar, R., B. Kaplan, N. Bouché, T. Arazi, D. Dolev, I. N. Talke, F. J. Maathuis, D. Sanders, D. Bouchez, and H. Fromm. 2000. “Expression of a Truncated Tobacco NtCBP4 Channel in Transgenic Plants and Disruption of the Homologous Arabidopsis CNGC1 Gene Confer Pb2+ Tolerance.” The Plant Journal: For Cell and Molecular Biology 24 (4): 533–42.

Sun, Sheng-Kai, Yi Chen, Jing Che, Noriyuki Konishi, Zhong Tang, Anthony J. Miller, Jian Feng Ma, and Fang-Jie Zhao. 2018. “Decreasing Arsenic Accumulation in Rice by Overexpressing OsNIP1;1 and OsNIP3;3 through Disrupting Arsenite Radial Transport in Roots.” The New Phytologist 219 (2): 641–53.

Takahashi, Ryuichi, Yasuhiro Ishimaru, Hugo Shimo, Yuko Ogo, Takeshi Senoura, Naoko K. Nishizawa, and Hiromi Nakanishi. 2012. “The OsHMA2 Transporter Is Involved in Root-to-Shoot Translocation of Zn and Cd in Rice.” Plant, Cell & Environment 35 (11): 1948–57.

Takano, Junpei, Motoko Wada, Uwe Ludewig, Gabriel Schaaf, Nicolaus von Wirén, and Toru Fujiwara. 2006. “The Arabidopsis Major Intrinsic Protein NIP5;1 Is Essential for Efficient Boron Uptake and Plant Development under Boron Limitation.” The Plant Cell 18 (6): 1498–1509.

Takemoto, Yuma, Yuta Tsunemitsu, Miho Fujii-Kashino, Namiki Mitani-Ueno, Naoki Yamaji, Jian Feng Ma, Shin-Ichiro Kato, Kozo Iwasaki, and Daisei Ueno. 2017. “The Tonoplast-Localized Transporter MTP8.2 Contributes to Manganese Detoxification in the Shoots and Roots of Oryza Sativa L.” Plant & Cell Physiology 58 (9): 1573–82.

Tanaka, Mayuki, Ian S. Wallace, Junpei Takano, Daniel M. Roberts, and Toru Fujiwara. 2008. “NIP6;1 Is a Boric Acid Channel for Preferential Transport of Boron to Growing Shoot Tissues in Arabidopsis.” The Plant Cell 20 (10): 2860–75.

Tanaka, Nobuhiro, Sho Nishida, Takehiro Kamiya, and Toru Fujiwara. 2016. “Large-Scale Profiling of Brown Rice Ionome in an Ethyl Methanesulphonate-Mutagenized Hitomebore Population and Identification of High-and Low-Cadmium Lines.” Plant and Soil 407 (1-2): 109–17.

Tarantino, Delia, Piero Morandini, Leonor Ramirez, Carlo Soave, and Irene Murgia. 2011. “Identification of an Arabidopsis Mitoferrinlike Carrier Protein Involved in Fe Metabolism.” Plant Physiology and Biochemistry: PPB /Societe Francaise de Physiologie Vegetale 49 (5): 520–29.

Tejada-Jiménez, Manuel, Rosario Castro-Rodríguez, Igor Kryvoruchko, M. Mercedes Lucas, Michael Udvardi, Juan Imperial, and Manuel González-Guerrero. 2015. “Medicago Truncatula Natural Resistance-Associated Macrophage Protein1 Is Required for Iron Uptake by Rhizobia-Infected Nodule Cells.” Plant Physiology 168 (1): 258–72.

Tejada-Jiménez, Manuel, Patricia Gil-Díez, Javier León-Mediavilla, Jiangqi Wen, Kirankumar S. Mysore, Juan Imperial, and Manuel González-Guerrero. 2017. “Medicago Truncatula Molybdate Transporter Type 1 (MtMOT1.3) Is a Plasma Membrane Molybdenum Transporter Required for Nitrogenase Activity in Root Nodules under Molybdenum Deficiency.” The New Phytologist 216 (4): 1223–35.

The Gene Ontology Consortium. 2017. “Expansion of the Gene Ontology Knowledgebase and Resources.” Nucleic Acids Research 45 (D1): D331–38.

Tian, Hui, Ivan R. Baxter, Brett Lahner, Anke Reinders, David E. Salt, and John M. Ward. 2010. “Arabidopsis NPCC6/NaKR1 Is a Phloem Mobile Metal Binding Protein Necessary for Phloem Function and Root Meristem Maintenance.” The Plant Cell 22 (12): 3963–79.

Ueno, Daisei, Akimasa Sasaki, Naoki Yamaji, Takaaki Miyaji, Yumi Fujii, Yuma Takemoto, Sawako Moriyama, et al. 2015. “A Polarly Localized Transporter for Efficient Manganese Uptake in Rice.” Nature Plants 1 (November): 15170.

Uraguchi, Shimpei, Nobuhiro Tanaka, Christian Hofmann, Kaho Abiko, Naoko Ohkama-Ohtsu, Michael Weber, Takehiro Kamiya, et al. 2017. “Phytochelatin Synthase Has Contrasting Effects on Cadmium and Arsenic Accumulation in Rice Grains.” Plant & Cell Physiology 58 (10): 1730–42.

Van Hoewyk, Douglas, Gulnara F. Garifullina, Ashley R. Ackley, Salah E. Abdel-Ghany, Matthew A. Marcus, Sirine Fakra, Keiki Ishiyama, et al. 2005. “Overexpression of AtCpNifS Enhances Selenium Tolerance and Accumulation in Arabidopsis.” Plant Physiology 139 (3): 1518–28.

Vitart, Veronique, Ivan Baxter, Peter Doerner, and Jeffrey F. Harper. 2001. “Evidence for a Role in Growth and Salt Resistance of a Plasma Membrane H+-ATPase in the Root Endodermis: Salt Sensitive H+-ATPase Mutant.” The Plant Journal: For Cell and Molecular Biology 27 (3): 191–201.

Von Wiren, N., S. Mori, H. Marschner, and V. Romheld. 1994. “Iron Inefficiency in Maize Mutant ys1 (Zea Mays L. Cv Yellow-Stripe) Is Caused by a Defect in Uptake of Iron Phytosiderophores.” Plant Physiology 106 (1): 71–77.

Wang, Chuang, Shan Ying, Hongjie Huang, Kuan Li, Ping Wu, and Huixia Shou. 2009. “Involvement of OsSPX1 in Phosphate Homeostasis in Rice.” The Plant Journal: For Cell and Molecular Biology 57 (5): 895–904.

Wang, Jing, Jinghan Sun, Jun Miao, Jinkao Guo, Zhanliang Shi, Mingqi He, Yu Chen, et al. 2013. “A Phosphate Starvation Response Regulator Ta-PHR1 Is Involved in Phosphate Signalling and Increases Grain Yield in Wheat.” Annals of Botany 111 (6): 1139–53.

Wang, Lu, Yinghui Ying, Reena Narsai, Lingxiao Ye, Luqing Zheng, Jingluan Tian, James Whelan, and Huixia Shou. 2013. “Identification of OsbHLH133 as a Regulator of Iron Distribution between Roots and Shoots in Oryza Sativa.” Plant, Cell & Environment 36 (1): 224–36.

Waters, Brian M., Heng-Hsuan Chu, Raymond J. Didonato, Louis A. Roberts, Robynn B. Eisley, Brett Lahner, David E. Salt, and Elsbeth L. Walker. 2006. “Mutations in Arabidopsis Yellow Stripe-like1 and Yellow Stripe-like3 Reveal Their Roles in Metal Ion Homeostasis and Loading of Metal Ions in Seeds.” Plant Physiology 141 (4): 1446–58.

Wild, Michael, Jean-Michel Davière, Thomas Regnault, Lali Sakvarelidze-Achard, Esther Carrera, Isabel Lopez Diaz, Anne Cayrel, Guillaume Dubeaux, Grégory Vert, and Patrick Achard. 2016. “Tissue-Specific Regulation of Gibberellin Signaling Fine-Tunes Arabidopsis Iron-Deficiency Responses.” Developmental Cell 37 (2): 190–200.

Wimalanathan, Kokulapalan, Iddo Friedberg, Carson M. Andorf, and Carolyn J. Lawrence-Dill. 2018. “Maize GO Annotation— Methods, Evaluation, and Review (maize-GAMER).” Plant Direct 2 (4): e00052.

Xu, Jiang, Hao-Dong Li, Li-Qing Chen, Yi Wang, Li-Li Liu, Liu He, and Wei-Hua Wu. 2006. “A Protein Kinase, Interacting with Two Calcineurin B-like Proteins, Regulates K+ Transporter AKT1 in Arabidopsis.” Cell 125 (7): 1347–60.

Xu, Jiming, Shulin Shi, Lei Wang, Zhong Tang, Tingting Lv, Xinlu Zhu, Xiaomeng Ding, Yifeng Wang, Fang-Jie Zhao, and Zhongchang Wu. 2017. “OsHAC4 Is Critical for Arsenate Tolerance and Regulates Arsenic Accumulation in Rice.” The New Phytologist 215 (3): 1090–1101.

Xu, Wenzhong, Wentao Dai, Huili Yan, Sheng Li, Hongling Shen, Yanshan Chen, Hua Xu, Yangyang Sun, Zhenyan He, and Mi Ma. 2015. “Arabidopsis NIP3;1 Plays an Important Role in Arsenic Uptake and Root-to-Shoot Translocation under Arsenite Stress Conditions.” Molecular Plant 8 (5): 722–33.

Yamaji, Naoki, Yuma Takemoto, Takaaki Miyaji, Namiki Mitani-Ueno, Kaoru T. Yoshida, and Jian Feng Ma. 2017. “Reducing Phosphorus Accumulation in Rice Grains with an Impaired Transporter in the Node.” Nature 541 (7635): 92–95.

Yang, An, Yansu Li, Yunyuan Xu, and Wen-Hao Zhang. 2013. “A Receptor-like Protein RMC Is Involved in Regulation of Iron Acquisition in Rice.” Journal of Experimental Botany 64 (16): 5009–20.

Yang, An, and Wen-Hao Zhang. 2016. “A Small GTPase, OsRab6a, Is Involved in the Regulation of Iron Homeostasis in Rice.” Plant & Cell Physiology 57 (6): 1271–80.

Yan, Jiali, Peitong Wang, Peng Wang, Meng Yang, Xingming Lian, Zhong Tang, Chao-Feng Huang, David E. Salt, and Fang Jie Zhao. 2016. “A Loss-of-Function Allele of OsHMA3 Associated with High Cadmium Accumulation in Shoots and Grain of Japonica Rice Cultivars.” Plant, Cell & Environment 39 (9): 1941–54.

Yan, Jiapei, Ju-Chen Chia, Huajin Sheng, Ha-Il Jung, Tetiana-Olena Zavodna, Lu Zhang, Rong Huang, et al. 2017. “Arabidopsis Pollen Fertility Requires the Transcription Factors CITF1 and SPL7 That Regulate Copper Delivery to Anthers and Jasmonic Acid Synthesis.” The Plant Cell 29 (12): 3012–29.

Yuan, Youxi, Huilan Wu, Ning Wang, Jie Li, Weina Zhao, Juan Du, Daowen Wang, and Hong-Qing Ling. 2008. “FIT Interacts with AtbHLH38 and AtbHLH39 in Regulating Iron Uptake Gene Expression for Iron Homeostasis in Arabidopsis.” Cell Research 18 (3): 385–97.

Zhai, Zhiyang, Sheena R. Gayomba, Ha-Il Jung, Nanditha K. Vimalakumari, Miguel Piñeros, Eric Craft, Michael A. Rutzke, et al. 2014. “OPT3 Is a Phloem-Specific Iron Transporter That Is Essential for Systemic Iron Signaling and Redistribution of Iron and Cadmium in Arabidopsis.” The Plant Cell 26 (5): 2249–64.

Zhang, Huimin, Yang Li, Xiani Yao, Gang Liang, and Diqiu Yu. 2017. “POSITIVE REGULATOR OF IRON HOMEOSTASIS1, OsPRI1, Facilitates Iron Homeostasis.” Plant Physiology 175 (1): 543–54.

Zhang, Lianhe, Bin Hu, Wei Li, Ronghui Che, Kun Deng, Hua Li, Feiyan Yu, Hongqing Ling, Youjun Li, and Chengcai Chu. 2014. “OsPT2, a Phosphate Transporter, Is Involved in the Active Uptake of Selenite in Rice.” The New Phytologist 201 (4): 1183–91.

Zhang, Ming, Yibo Cao, Zhiping Wang, Zhi-Qiang Wang, Junpeng Shi, Xiaoyan Liang, Weibin Song, Qijun Chen, Jinsheng Lai, and Caifu Jiang. 2018. “A Retrotransposon in an HKT1 Family Sodium Transporter Causes Variation of Leaf Na+ Exclusion and Salt Tolerance in Maize.” The New Phytologist 217 (3): 1161–76.

Zhang, Yuanyuan, Kai Chen, Fang-Jie Zhao, Cuiju Sun, Cheng Jin, Yuheng Shi, Yangyang Sun, et al. 2018. “OsATX1 Interacts with Heavy Metal P1B-Type ATPases and Affects Copper Transport and Distribution.” Plant Physiology 178 (1): 329–44.

Zhang, Yu, Yong-Han Xu, Hong-Yin Yi, and Ji-Ming Gong. 2012. “Vacuolar Membrane Transporters OsVIT1 and OsVIT2 Modulate Iron Translocation between Flag Leaves and Seeds in Rice.” The Plant Journal: For Cell and Molecular Biology 72 (3): 400–410.

Zhao, Meng, Hong Ding, Jian-Kang Zhu, Fusuo Zhang, and Wen-Xue Li. 2011. “Involvement of miR169 in the Nitrogen-Starvation Responses in Arabidopsis.” The New Phytologist 190 (4): 906–15.

Zhao, Xue Qiang, Namiki Mitani, Naoki Yamaji, Ren Fang Shen, and Jian Feng Ma. 2010. “Involvement of Silicon Influx Transporter OsNIP2;1 in Selenite Uptake in Rice.” Plant Physiology 153 (4): 1871–77.

Zheng, Luqing, Naoki Yamaji, Kengo Yokosho, and Jian Feng Ma. 2012. “YSL16 Is a Phloem-Localized Transporter of the Copper-Nicotianamine Complex That Is Responsible for Copper Distribution in Rice.” The Plant Cell 24 (9): 3767–82.

Zhou, Jie, Fangchang Jiao, Zhongchang Wu, Yiyi Li, Xuming Wang, Xiaowei He, Weiqi Zhong, and Ping Wu. 2008. “OsPHR2 Is Involved in Phosphate-Starvation Signaling and Excessive Phosphate Accumulation in Shoots of Plants.” Plant Physiology 146 (4): 1673–86.

Zhu, Jinsheng, Kelvin Lau, Robert Puschmann, Robert K. Harmel, Youjun Zhang, Verena Pries, Philipp Gaugler, et al. 2019. “Two Bifunctional Inositol Pyrophosphate Kinases/phosphatases Control Plant Phosphate Homeostasis.” eLife 8 (August). https://doi.org/10.7554/eLife.43582.

Zimeri, Anne Marie, Om Parkash Dhankher, Bonnie McCaig, and Richard B. Meagher. 2005. “The Plant MT1 Metallothioneins Are Stabilized by Binding Cadmiums and Are Required for Cadmium Tolerance and Accumulation.” Plant Molecular Biology 58 (6): 839–55.

